# Inhibition of CDK12 elevates cancer cell dependence on P-TEFb by stimulation of RNA polymerase II pause release

**DOI:** 10.1101/2023.03.21.533608

**Authors:** Zhijia Wang, Samu V. Himanen, Heidi M. Haikala, Caroline C. Friedel, Anniina Vihervaara, Matjaž Barborič

## Abstract

P-TEFb and CDK12 facilitate transcriptional elongation by RNA polymerase II. Given the prominence of both kinases in cancer, gaining a better understanding of their interplay could inform the design of novel anti-cancer strategies. While down-regulation of DNA repair genes in CDK12-targeted cancer cells is being explored therapeutically, little is known about mechanisms and significance of transcriptional induction upon inhibition of CDK12. We show that selective targeting of CDK12 in colon cancer-derived cells activates P-TEFb via its release from the inhibitory 7SK snRNP. In turn, P-TEFb stimulates Pol II pause release at thousands of genes, most of which become newly dependent on P-TEFb. Amongst the induced genes are those stimulated by hallmark pathways in cancer, including p53 and NF-κB. Consequently, CDK12-inhibited cancer cells exhibit hypersensitivity to inhibitors of P-TEFb. While blocking P-TEFb triggers their apoptosis in a p53-dependent manner, it impedes cell proliferation irrespective of p53 by preventing induction of genes downstream of the DNA damage-induced NF-κB signaling. In summary, stimulation of Pol II pause release at the signal-responsive genes underlies the functional dependence of CDK12-inhibited cancer cells on P-TEFb. Our study establishes the mechanistic underpinning for combinatorial targeting of CDK12 with either P-TEFb or the induced oncogenic pathways in cancer.

## INTRODUCTION

Transcriptional cyclin-dependent kinases (tCDKs) phosphorylate Tyr^1^Ser^2^Pro^3^Thr^4^Ser^5^Pro^6^Ser^7^ heptad repeats within the C-terminal domain (CTD) of the RPB1 subunit of RNA polymerase II (Pol II) and Pol II accessory factors. This promotes transitions between discrete phases of the transcription cycle and links transcription with precursor mRNA (pre-mRNA) maturation and chromatin modification (1-3). Transcription initiation and promoter clearance is stimulated by CDK7. As a part of the ten-subunit transcription factor TFIIH, CDK7 binds Cyclin H and Mat1 and is recruited to the Pol II pre-initiation complex at gene promoters where it phosphorylates Ser5 and Ser7 residues of the Pol II CTD. At the same time, CDK7 promotes 5’-end capping of pre-mRNA and facilitates promoter-proximal pausing of Pol II by stimulating the binding of DRB-sensitivity inducing factor (DSIF) and negative elongation factor (NELF) with Pol II, which leads to its arrest in an inactive conformation (1,4). Outside of TFIIH, the trimeric CDK7-cyclin H-Mat1 complex functions as a CDK-activating kinase (CAK) for the Pol II elongation kinases positive transcription elongation factor b (P-TEFb), CDK12 and CDK13 (5). The release of Pol II from pausing is triggered by the recruitment of P-TEFb, composed of the catalytic CDK9 and a regulatory cyclin T1 or T2 subunit, which phosphorylates Ser2 residues across the CTD and its linker as well as multiple positions within both pause-inducing factors (6,7). These events convert DSIF into a positive elongation factor, lead to dissociation of NELF from Pol II, and stimulate the docking of the polymerase associated factor complex (PAFc) and SPT6 onto Pol II and the CTD linker, respectively, enabling Pol II elongation to resume (8). Subsequently, highly homologous CDK12 and CDK13 interact with cyclin K to phosphorylate the CTD at multiple positions and to stimulate processivity and elongation rate of Pol II along gene bodies by promoting the interaction of PAFc and SPT6 with elongating Pol II, which facilitates production of full-length transcripts (9-13). In keeping with their role during transcription, both P-TEFb and CDK12 regulate alternative splicing of pre-mRNA and proper transcription termination (14-19).

CDK12 has emerged as an important player in cancer. An analysis of the genetic dependency data across 1070 cancer cell lines from the Cancer Dependency Map (DepMap) project has revealed that *CDK12* is essential in close to 30% of cancer cell lines, of which average knockout effects on cell viability were similar to those of the cell-cycle oncogenes *CDK4* and *CDK6* (20). Genomic alterations of *CDK12*, which include mutations, deletions, amplifications and rearrangements, were documented in about thirty tumor types with an incidence of up to 15% of cases (21). The best understood role of CDK12 is its ability to enable genomic stability, where CDK12 promotes expression of unique sets of genes. First, CDK12 facilitates proper expression of long genes that function in the homologous recombination (HR) DNA repair pathway, DNA replication and cell-cycle. Therein, its activity counteracts a prevalence of intronic polyadenylation sites to suppress premature pre-mRNA cleavage and polyadenylation (PCPA) (9,10,22,23). Second, CDK12 stimulates translation of mRNAs encoding key factors that dictate chromosome alignment and mitotic fidelity (24). Whereas disruption of these CDK12 regulatory networks likely contributes to the origin of tandem DNA duplications that are a hallmark of genomic instability in cancers with biallelic *CDK12* inactivation (23,25,26), the requirement for HR gene expression has invigorated efforts for therapeutic targeting of CDK12, particularly in combination with PARP inhibitors that are effective in HR-deficient cancer cells (27-29). As outcomes of *CDK12* alterations are context dependent, *CDK12* has been reported to function as a tumor suppressor and as an oncogene. For example, *CDK12* is a tumor suppressor in high-grade serous ovarian carcinoma, wherein its recurrent loss-of-function mutations prompt downregulation of HR genes, impairment of DNA double-strand break (DSB) repair via HR and cellular sensitivity to PARP inhibitors (30-32). Conversely, *CDK12* is co-amplified with the receptor tyrosine kinase (RTK) oncogene *HER2* (also known as *ERBB2*) in many cases of breast cancer, where it promotes migration and invasion of tumor cells by downregulation of the nuclear isoform of DNAJB6 via alternative last exon splicing (15). In addition, CDK12 can act as a facilitator or an effector of oncogenic signaling pathways. While CDK12 facilitates activation of ERBB–PI3K–AKT and WNT signaling pathways in *HER2*-positive breast cancer through promoting transcription of an RTK adaptor IRS-1 and WNT ligands genes, respectively (33), it is constitutively activated by the hyperactive RAS–BRAF–MEK1/2–ERK1/2 signaling pathway in BRAF-mutated melanoma (34).

In contrast to CDK12, current evidence indicates that the role of P-TEFb in cancer is chiefly, if not exclusively, oncogenic. An analysis of the DepMap dataset shows that *CDK9*, just like *CDK7*, is essential for all cancer cell lines, of which knockout displays even stronger average effects than the knockout of *KRAS* or the cell-cycle *CDK4* and *CDK6* (20). Because cancer alterations of *CDK9* do not affect its expression or kinase activity in a majority of cases (35,36), many distinct mechanisms likely account for the reliance on P-TEFb. For example, genetic and epigenetic alterations in cancer cells, most notably rearrangements of *MLL1* and amplification of *MYC*, respectively, lead to redirection of P-TEFb to the oncogenes’ loci, stimulating oncogenic transcription programs (37). Furthermore, P-TEFb promotes enhancer-driven transcription of *MYC* and *MCL1* oncogenes and protein stability of MYC to underwrite cancer cell dependency on these short-lived oncogenic proteins (38-40). P-TEFb also stimulates transcription of key transcription factors (TFs) of epithelial–mesenchymal transition (EMT) to promote breast cancer EMT, invasion and metastasis (41). Recently, P-TEFb has been shown to suppress mitochondria-mediated intrinsic apoptosis pathway via facilitating transcription of key genes of the oncogenic PI3K–AKT signaling cascade (42). Beyond its role in Pol II pause release, P-TEFb also promotes tumorigenesis by silencing tumor suppressor and endogenous retrovirus genes through phosphorylation of an ATP-dependent helicase BRG1 of the BAF chromatin remodeling complex, which is thought to keep this chromatin remodeler off chromatin (43).

Transcriptional activity of P-TEFb is controlled by the 7SK small nuclear ribonucleoprotein (snRNP), in which coordinated actions of the scaffolding 7SK small nuclear RNA (7SK) and three RNA-binding proteins (RBPs) inhibit the kinase. While the 7SK γ-methylphosphate capping enzyme MePCE and La-related protein 7 (LARP7) stabilize 7SK to form the tripartite core of 7SK snRNP, HEXIM1 interacts with 7SK of the core to bind and inhibit P-TEFb (44,45). As 7SK snRNP contains a major fraction of cellular P-TEFb, it acts as a molecular reservoir from which active P-TEFb can be deployed in a prompt and regulated manner to accommodate cellular transcriptional needs in unison with stimuli-specific transcription factors that recruit the released P-TEFb to promote target gene activation. Indeed, a host of perturbations and stimuli, including transcriptional blockade, genotoxic stress, viral infection and cell growth, trigger activation of P-TEFb, whereby numerous 7SK-binding factors have been proposed to release P-TEFb enzymatically and/or allosterically via a large-scale conformational switching of 7SK (45,46).

While CDK12-dependent transcription has been explored extensively and harnessed in therapeutic settings, little attention has been given to genes that are induced upon the inhibition of CDK12. In principle, the induced genes could elicit novel vulnerabilities and/or resistance mechanisms of CDK12- targeted cancer cells. Intriguingly, transcriptional responses to CDK12 inhibition and ultraviolet (UV) irradiation resemble each other in that they both show a gene length-dependent Pol II elongation effect, wherein transcription of short genes is induced or remains unaltered, whereas increasingly long genes become progressively repressed (10,47,48). Importantly, while prolonged perturbation of CDK12 increases DNA damage (9,27,28,31,32), the release of P-TEFb from 7SK snRNP is a hallmark of cellular DNA damage response (DDR), whereby activated P-TEFb shapes a transcriptional response enabling survival of stressed cells (49,50). In this study, we investigated whether inhibition of CDK12 induces genes via activation of P-TEFb. Furthermore, we explored the nature and functional significance of the induced transcriptome. Our results show that inhibition of CDK12 stimulates P-TEFb and induces genes downstream of important cancer signaling pathways, which renders cancer cells highly sensitive to P-TEFb inhibitors. Hence, our study provides a rationale for elimination of cancer cells by co-targeting the principal kinases that stimulate transcription elongation by Pol II.

## MATERIALS AND METHODS

### Reagents

A list of reagents used in the study is provided in Supplementary Table S1.

### Cell lines

HCT116 cell line and its derivatives were cultured in McCoy’s 5A Medium (Merck) supplemented with 10% FBS and 100 U/ml penicillin/streptomycin. CCD 841 CoN cell line was cultured in Eagle’s Minimum Essential Medium (Lonza) supplemented with 10% FBS and 100 U/ml penicillin/streptomycin. All cell lines were maintained at 37°C with 5% CO_2_. Cell lines were confirmed regularly to be mycoplasma-free using a qPCR-based Mycoplasmacheck detection kit (Eurofins). Cell lines were not authenticated by us, but retrieved from trusted sources as listed in Supplementary Table S1D.

### Chemicals

NVP-2, THZ531 and MLN120B were from MedChemExpress. THZ532 and YLK-5-124 were a gift from Nathanael S. Gray’s Laboratory (Stanford University, USA). iCDK9 was a gift from Qiang Zhou’s Laboratory (University of California, Berkley, USA). 3-MB-PP1 was from Cayman Chemical. KU-60019 was from Selleck Chemicals. TNF-α was from Proteintech. For more information on the chemicals, see Supplementary Table S1B.

### Co-immunoprecipitation Assay

HCT116 and HCT116 *CDK12* ^as/as^ cells grown on 60 mm plates were treated as indicated at 90 % confluency, after which whole-cell extracts were prepared using 800 μl of lysis buffer C (20 mM Tris-HCl, 0.5% NP-40, 150 mM NaCl, 1.5 mM MgCl2, 10 mM KCl, 10% Glycerol, 0.5 mM EDTA, pH 7.9) on ice for 1 h in the presence of EDTA-free Protease Inhibitor Cocktail (Merck) and cleared by centrifugation at 20,000 g for 15 min. Buffer C-equilibrated Dynabeads Protein G (Thermo Fisher Scientific) were incubated with 1 μg of the anti-HEXIM1 antibody (Everest Biotech) for 2 hr at 4 °C and washed three times with the lysis buffer prior to incubation with 800 μl of the whole-cell extracts overnight at 4 °C. The beads were washed three times with the lysis buffer and boiled in SDS loading buffer supplied with 10% of β-Mercaptoethanol for 5 min. Proteins were separated using SDS-PAGE, transferred to nitrocellulose membrane and analyzed by Western blotting. Thirty μl of the whole-cell extract was used as a loading control.

### Western Blotting Assay and Antibodies

Whole-cell extracts were prepared using lysis buffer C (20 mM Tris-HCl, 0.5% NP-40, 150 mM NaCl, 1.5 mM MgCl2, 10 mM KCl, 10% Glycerol, 0.5 mM EDTA, pH 7.9) on ice for 1 h in the presence of EDTA-free Protease Inhibitor Cocktail (Merck). Lysates were cleared by centrifugation at 20,000 g for 15 min, boiled in SDS loading buffer supplied with 10% of β-Mercaptoethanol for 5 min, separated using SDS-PAGE, transferred to nitrocellulose membrane and analyzed by Western blotting. The following antibodies were used according to manufacturers’ instructions: anti-CDK9 (Santa Cruz Biotechnology, 1:4000); anti-p53 (Santa Cruz Biotechnology, 1:3000); anti-p21 (Santa Cruz Biotechnology, 1:4000); anti-GAPDH (Santa Cruz Biotechnology, 1:10,000); anti-HEXIM1 (Everest Biotech, 1:000); anti-CDK12 (Cell Signaling Technology, 1:1000); anti-CDK7 (Cell Signaling Technology, 1:1000). Manufacturers provide validation for all antibodies. For more information on the antibodies, see Supplementary Table S1A.

### Target Engagement Assay

HCT116 cells were treated with the indicated concentrations of THZ531 or DMSO for the indicated time. Whole-cell extracts were prepared using NP-40 lysis buffer (Thermo Fisher Scientific) supplemented with complete protease inhibitor cocktail (Merck), PhosSTOP phosphatase inhibitor cocktail (Merck) and 1 mM of Phenylmethanesulfonyl fluoride (Merck). The extracts containing 1 mg of total protein were first incubated with 1 µM of biotin-THZ1 at 4 °C overnight, after which the complexes were affinity purified using 30 µl of streptavidin agarose beads for 2 h at 4 °C. The beads were washed three times with the lysis buffer and boiled for 10 min in SDS loading buffer. The proteins were separated using SDS-PAGE and analyzed by Western blotting. Fifty micrograms of total protein was used as a loading control.

### RNA Extraction and RT-qPCR Assay

HCT116 cells were seeded on 6-well plates for 16 h and treated at 80% confluency with the indicated compound combinations and corresponding volume of DMSO. RNA samples were extracted using TRI Reagent (Merck), DNase-treated with the Turbo DNA-Free kit (Thermo Fisher Scientific), and reverse transcribed with M-MLV reverse transcriptase (Thermo Fisher Scientific) and random hexamers (Thermo Fisher Scientific) according to the manufacturers’ instructions. qPCR reactions were performed with diluted cDNAs, primer pairs that spanned exon-intron junctions, and FastStart Universal SYBR Green QPCR Master (Rox) (Sigma) using Roche LightCycler 480. Primers were from Integrated DNA Technologies and designed using PrimerQuest Tool. Results from three independent experiments were normalized to the values of *GAPDH* mRNA and DMSO-treated cells, and plotted as the mean ± s.e.m. Sequences of the primers are listed in Supplementary Table 1F.

### Cytotoxicity, Viability and Apoptosis Assays

Cytotoxicity, viability, and apoptosis assays were conducted using CellTox Green Cytotoxicity Assay (Promega), CellTiter-Glo 2.0 Cell Viability Assay (Promega) and RealTime-Glo Annexin V Apoptosis and Necrosis Assay (Promega), respectively, according to manufacturer’s instructions. For cytotoxicity and apoptosis assays, HCT116 cells were seeded at density of 20,000 cells/well on 96-well plates for 16 h prior to the treatments. For the seven-day viability assays, HCT116 cells were seeded at density of 5,000 cells/well on 12-well plates for 16 h prior to the treatments. Cell culture medium and drugs were replenished on day 4 of the assay. Fluorescence and luminescence measurements were taken at the indicated time points using PerkinElmer Victor X3 reader. Results from at least three independent experiments are presented as values relative to the values of DMSO-treated cells and plotted as the mean ± s.e.m.

### Cell Confluency assay

The assay was performed using CELLCYTE X™ live cell imager and analyzer (Cytena). The parental and *TP53* ^−/−^ HCT116 cells were seeded at density of 1,000 cells/well and 1,250 cells/well, respectively, on 96-well plates for 16 h prior to the treatments. Cells were imaged every four hours for seven days. Cell culture medium and drugs were replenished on day 4 of the assay. GraphPad Prism v. 9.5.1 was used for plotting and statistical analysis of the results from three independent experiments.

### Spheroid Apoptosis Assay

HCT116 cells were seeded at density of 400 cells/well on Nunclon Sphera 96-well U-bottom ultra low attachment 3D cell culture plate (Thermo Fisher Scientific) and spun down at 200 g for 10 minutes. After culturing the cells for 48 h to allow for spheroid formation, the spheroids were treated with the indicated compounds over the subsequent 48 h period. Apoptosis assay was conducted using RealTime-Glo Annexin V Apoptosis and Necrosis Assay (Promega) according to manufacturer’s instructions. Luminescence measurements were taken at the indicated time points using PerkinElmer Victor X3 reader. Results from at least three independent experiments are presented as values relative to the values of DMSO-treated cells and plotted as the mean ± s.e.m.

### Cell-cycle analysis

HCT116 cells were seeded on 60 mm dishes and cultured to approximately 70% confluence. The cells were synchronized by serum deprivation for 24 h prior to their culture in the serum-containing medium for 24 h in the presence of DMSO (-), THZ531 (50 nM) and NVP-2 (50 nM). After trypsinization, cell pellets were collected, washed with PBS and fixed by ice-cold 70% ethanol. For FACS analysis, the cells were washed in PBS and incubated overnight at 4°C in freshly prepared staining solution in PBS containing 50 g/ml propidium iodide, 0.1% Triton X-100 and 0.2 mg/ml DNase-free Rnase A. FACS was performed by BD Accuri C6 and data was analysed by FlowJo.

### Analysis of RNA Sequencing Datasets

Log2 fold-changes in gene expression and multiple testing adjusted *P*-values for the RNA sequencing datasets listed below were taken from the corresponding studies. No re-analysis of the raw RNA-seq data was performed, except for the data from the study of Houles *et al.* (34) (see the details below). Gene expression changes in nuclear RNA obtained from HCT116 *CDK12* ^as/as^ cells treated with 3-MB-PP1 for 4.5 h were taken from the study of Manavalan *et al.* (Dataset EV2 from this publication) (23). The RNA-seq data are available at the Gene Expression Omnibus (GEO) repository (NCBI) under the accession code GSE120071. Gene expression changes obtained from A-375 and Colo829 cells treated with THZ531 (500 nM) for 6 h were taken from the study of Houles *et al.* (34). As Houles *et al.* did not specify how log2 fold-changes were calculated and whether *P*-values were adjusted for multiple testing, we re-calculated log2 fold-changes and adjusted *P*-values using DESeq2 (51). For this purpose, we downloaded raw read count matrices for all genes from the GEO repository using the accession code GSE184734, and used this as the input for DESeq2. Gene expression changes obtained from MDA-MB-231 cells treated with SR-4835 (90 nM) for 6 h were taken from the study of Quereda *et al.* (28) (Table S4 from this publication). The RNA-seq are available at the BioProject database (NCBI) under the accession code PRJNA530774. TT-seq expression changes obtained from IMR-32 cells treated with THZ531 (400 nM) for 2 h were taken from the study of Krajewska *et al.* (Supplementary Data 2 from this publication) (10). The TT-seq data are available at the GEO repository under the accession code GSE113313. In all of these studies, log2 fold-changes were determined either with DESeq2 or edgeR (52) and *P*-values were adjusted for multiple testing with the Benjamini and Hochberg method (53).

Genes were considered significantly regulated if the multiple testing adjusted *P*-value the was ≤ 0.001, up-regulated with a log2 fold-change ≥ 1 and down-regulated with a log2 fold-change ≤ -1. Gene lengths were obtained from Ensembl. Distribution of gene lengths for up-and down-regulated genes is shown as a boxplot, which was created in R (https://www.R-project.org/). Boxes represent the range between the first and third quartiles for each condition. Gray horizontal lines in boxes show the median. The ends of the whiskers (vertical lines) extend the box by 1.5 times the inter-quartile range. Data points outside this range (outliers) are shown as small circles. Gene set enrichment analysis on log2 fold-changes for protein-coding genes was performed using the command line tool from http://www.gsea-msigdb.org/ (version 4.0.2) (54) and hallmark gene sets from Molecular Signatures Database (55).

### Precision Run-On sequencing

PRO-seq and its data analyses were conducted as previously described (56,57), with minor modifications. HCT116 cells were seeded on 15 cm dishes and cultured for 14 h to approximately 90% confluence, reaching approximately 17 million cells per a dish. Subsequently, the cells were treated with DMSO, 3-MB-PP1 (5 µM) and NVP-2 (10 nM) for either 1 h or 4,5 h. Chromatin was isolated with NUN buffer (0.3 M NaCl, 7.5 mM MgCl_2_, 1M Urea, 1% NP-40, 20 mM HEPES pH 7.5, 0.2 mM EDTA,1 mM DTT, 20 U SUPERaseIn RNase Inhibitor [Life Technologies], 1 x EDTA-free Protease Inhibitor Cocktail [Roche]). The samples were centrifuged (12 500 g, 30 min, 4°C), after which the chromatin pellets were washed with 50 mM Tris-HCl (pH 7.5) and stored at -80°C in chromatin storage buffer (50mM Tris-HCl pH 8.0, 5 mM MgAc_2_, 25% glycerol, 0.1 mM EDTA, 5mM DTT, 20 units SUPERaseIn RNase Inhibitor). The isolated chromatin was fragmented by sonication at 4°C for 10 minutes using Bioruptor (Diagenode). Run-on reactions were performed at 37°C for 5 minutes using biotinylated nucleotides (5 mM Tris-HCl pH 8.0, 150 mM KCl, 2.5 mM MgCl_2_, 0.5 mM DTT, 0.5% Sarkosyl, 0.4 U/µl RNase inhibitor, 0.025 mM biotin-ATP/CTP/GTP/UTP [Perkin Elmer]). Total RNA was extracted using Trizol and ethanol precipitation. Isolated RNA was fragmented by base hydrolysis with NaOH. Nascent RNAs labeled with biotin were isolated from total RNA using streptavidin-coated beads (Dynabeads MyOne Streptavidin C1, Thermo Fisher Scientific). TruSeq small RNA adaptor was ligated to 3’ ends of nascent RNAs and transcripts purified with a second round of streptavidin bead purification. 5’-cap was removed using RNA 5′ pyrophosphohydrolase (Rpph, NEB) followed by addition of a phosphate group to the 5’ end using T4 polynucleotide kinase (T4 PNK, NEB). Next, Truseq small RNA adaptor was ligated to the repaired 5’ end of nascent RNAs, followed by a third round of purification with the streptavidin beads, and reverse transcription was used to convert adaptor-ligated RNAs to cDNA. Finally, PRO-seq libraries were amplified with PCR and sequenced with Illumina using paired-end, 50nt+50nt settings. Sequences of the adaptors and primers are listed in Supplementary Table 1G.

### PRO-seq data analyses

Adapters sequences in read 1 (TGGAATTCTCGGGTGCCAAGGAACTCCAGTCAC) and read 2 (GATCGTCGGACTGTAGAACTCTGAACGTGTAGATCTCGGTGGTCGCCGTATCATT) were removed, and PCR duplicates collapsed (based on 6-nt UMI in the 5’-adapter), using fastp tool (58). The reads were aligned to human genome version hg38 using Bowtie2 (59) in paired-end mode with - -end-to-end parameter. Regulatory element identification from global run-on sequencing data (dREG) (60) was used to identify actively transcribed genes displaying initiation of transcription at the transcription start site (TSS). Transcription at actively transcribed genes was quantified at promoter-proximal (–250 to +250 nucleotide from the TSS) and gene body regions (+250 nucleotide from the TSS to –500 nucleotide from the cleavage and polyadenylation site [CPS]). Differentially transcribed genes (DTGs) in response to treatments were determined from Pol II engagements at gene bodies using DESeq2 (51). Adjusted *P*-value ≤ 0.05, and fold-change ≥ 1.25 for induced and ≤ 0.8 for down-regulated genes were used. MA plots show average log2 of read counts at gene bodies on x-axis and log2 fold-changes on the y-axis.

### Visualizing engaged Pol II at genes

Density profiles of nascent transcription were generated as previously described (57), from the 3’-nts of normalized PRO-seq reads using bedtools genome coverage (61) and bedgraphtobigwig (62). In the genome browser, the scale of the y-axis is linear and equal to all samples. The signal on the plus strand is shown on a positive scale, and the signal on the minus strand is shown on a negative scale. The metagene profiles were generated from the normalized bigWig files using bigwig package (https://github.com/andrelmartins/bigWig/) using a query window of 10 nucleotides (around the TSS or CPS), or a 1/50^th^ fraction of the gene body. Pausing index (PI) was quantified as Pol II density at promoter-proximal region / Pol II density across the gene body. The treatment-induced change in the PI was quantified at every transcribed gene as PI upon the indicated tCDK inhibitor treatment – PI of the time-matched DMSO control.

### Quantification and Statistical Analysis

*P*-values of gene expression changes were corrected by multiple testing using the method by Benjamini and Hochberg (53) for adjusting the false discovery rate (FDR) and an adjusted *P*-value cutoff of 0.001 was applied. Data shown for all RT-qPCR experiments and functional assays were collected from three biological replicates as indicated in individual figure legends and are presented as means ± s.e.m. Statistical significance and *P*-values were determined by Student’s *t* test or two-way ANOVA between the indicated paired groups of biological replicates. Where the results were not statistically significant, *P*-values are not indicated.

## RESULTS

### Inhibition of CDK12 stimulates the release of P-TEFb from 7SK snRNP

Considering that perturbation of CDK12 increases DNA damage (9,27,28,31,32) and that genotoxic stress triggers activation of P-TEFb via its release from 7SK snRNP to enable a pro-survival transcriptional response (49,50), we hypothesized that selective inhibition of CDK12 activates P-TEFb (Figure 1A). To test this prediction, we performed co-immunoprecipitation (co-IP) assays to monitor the interaction of endogenous CDK9 with its inhibitory protein HEXIM1 in whole cell extracts of cells that were treated or not with inhibitors of CDK12. We conducted these experiments using HCT116 cells, a well-established model of colorectal cancer encoding the oncogenic KRAS ^G13D^ and PIK3CA ^H1047R^ proteins. We first inhibited CDK12 with THZ531, a potent and selective molecular probe which covalently modifies the cysteine residue 1039 positioned outside of the conserved kinase domain (63). Treatment of HCT116 cells with 300 or 400 nM of THZ531 resulted in a time-dependent dissociation of CDK9 from HEXIM1, indicative of P-TEFb activation (Figure 1B). In contrast, 400 nM of THZ532, the enantiomer of THZ531 with a diminished ability to inhibit CDK12 (63), failed to do so.

**Figure 1.**
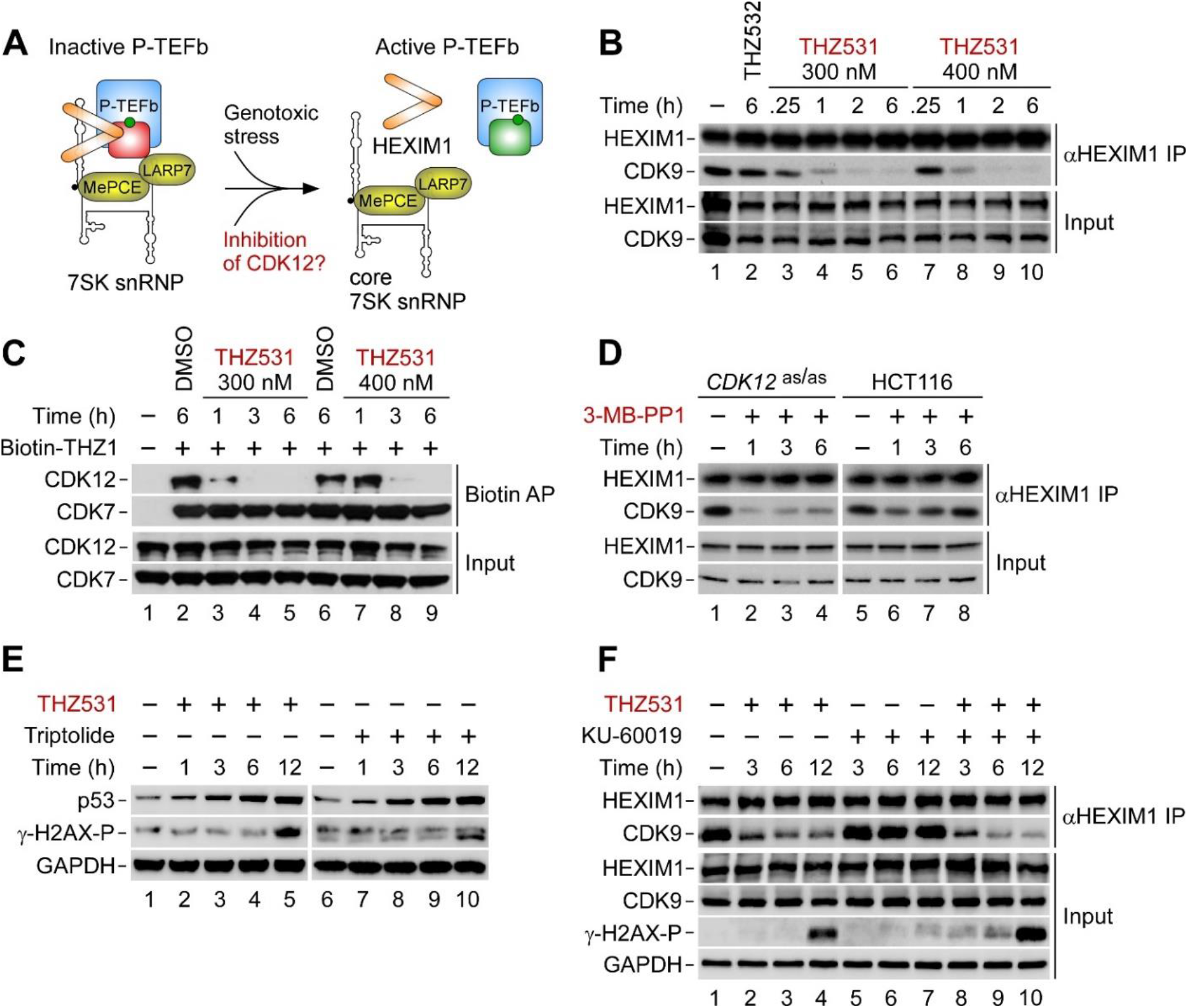
**Inhibition of CDK12 triggers the release of P-TEFb from the inhibitory 7SK snRNP complex.** (A) Schematic representation of controlling the kinase activity of P-TEFb with 7SK snRNP. Upon the release of P-TEFb from 7SK snRNP, the interaction of P-TEFb with HEXIM1 does not take place, resulting in the activation of P-TEFb. For simplicity, a large-scale conformational switching of 7SK upon the release of P-TEFb from the core of 7SK snRNP is not depicted. (B) CoIP of HEXIM1 with CDK9 from whole cell extracts of HCT116 cells. Cells were treated with DMSO (lane 1), THZ532 (400 nM) and two different doses of THZ531 for the indicated duration prior to preparation of whole cell extracts and detection of HEXIM1 and CDK9 in HEXIM1 immunoprecipitations (αHEXIM1 IP) and whole cell extracts (Input) by Western blotting. (C) Specificity of the targeting of CDK12 with THZ531 in target engagement assay. HCT116 cells were treated with DMSO and two different doses of THZ531 for the indicated duration (in hours) prior to preparation of whole cell extracts and detection of CDK12 and CDK7 in Biotin-THZ1 (1 μM) affinity purifications (Biotin AP) and whole cell extracts (Input) by Western blotting. (D) CoIP of HEXIM1 with CDK9 from whole cell extracts of the indicated HCT116 cell lines. Cells were treated with DMSO (lanes 1 and 5) and 3-MB-PP1 for the indicated duration (in hours) prior to preparation of whole cell extracts and detection of HEXIM1 and CDK9 in αHEXIM1 IP and whole cell extracts (Input) by Western blotting. (E) HCT116 cells were treated with DMSO (lanes 1 and 6), THZ531 (400 nM) and triptolide (1 μM) as indicated for the indicated duration (in hours) prior to preparation of whole cell extracts and detection of the indicated proteins by Western blotting. (F) CoIP of HEXIM1 with CDK9 from whole cell extracts of HCT116 cells. Cells were treated with DMSO (-), THZ531 (400 nM) and KU-60019 (5 μM) alone and in combination as indicated for the indicated duration prior to preparation of whole cell extracts and detection of HEXIM1 and CDK9 in αHEXIM1 IP and the indicated proteins in whole cell extracts (Input) by Western blotting.

Two lines of evidence indicate the on-target effect of THZ531 in our system. First, both concentrations of THZ531 but not 400 nM of THZ532 led to a time-and dose-dependent decrease in mRNA levels of *BRCA1* and *FANCD2*, the established CDK12-dependent genes of the HR pathway (9,31) (Supplementary Figure S1A). Second, we conducted target engagement assay to assess whether the THZ531 concentrations targeted CDK12 specifically over CDK7. Namely, THZ531 displays a residual, more than 50-fold lower activity towards CDK7 and CDK9 in fixed-endpoint *in vitro* kinase assays (63). We used a biotin-modified version of THZ1, a covalent inhibitor of CDK7 that interacts with both CDK7 and CDK12 (63,64), to affinity purify both kinases from HCT116 cell extracts and assess whether THZ531 prevents CDK12 but not CDK7 from binding the biotinylated THZ1. Indeed, treatment of HCT116 cells with 300 nM or 400 nM of THZ531 countered the affinity purification of CDK12 in time-dependent manner, while CDK7 remained bound to biotinylated THZ1 across all conditions tested (Figure 1C).

However, in addition to CDK12, THZ531 shows a comparable potency towards the homologous CDK13 (63), which is not involved in facilitating DNA repair (9,65). To demonstrate that inhibition of CDK12 alone provokes the release of CDK9 from HEXIM1, we took a chemical genetic approach. We conducted co-IP assays in HCT116 *CDK12* ^as/as^ cell line in which both alleles of *CDK12* were replaced with *CDK12* ^F813G^, generating analog-sensitive version of CDK12 (asCDK12) that is specifically inhibited by treatment of these cells with bulky adenine analogs such as 3-MB-PP1 (23). Recapitulating the results with THZ531, treatment of HCT116 *CDK12* ^as/as^ cell line with 3-MB-PP1 perturbed the interaction of CDK9 with HEXIM1 (Figure 1D). In contrast, we did not observe this effect in the parental HCT116 cell line which is refractory to the inhibition of CDK12 by 3-MB-PP1 (Figure 1D). Confirming that 3-MB-PP1 specifically inhibited asCDK12, 3-MB-PP1 treatment decreased mRNA levels of *BRCA1* and *FANCD2* in HCT116 *CDK12* ^as/as^ cells but not in the parental counterpart (Supplementary Figure S1B).

Given that perturbation of CDK12 induces DSBs (9,27,28,31,32), we asked whether elevated levels of DSBs stimulate the release of P-TEFb from 7SK snRNP in CDK12-inhibited cells. Here, we followed the levels of DSBs-induced Ser139 phosphorylation of a histone variant γ-H2AX (γ-H2AX-P) in cell extracts of HCT116 cells treated with 400 nM of THZ531 using immunoblotting analysis. We detected a robust increase in the levels of γ-H2AX-P at twelve hours into the treatment but not during the first six hours when CDK9 dissociates from HEXIM1 (Figure 1E, left). Within this initial period, however, we detected accumulation of p53, suggesting that transcriptional blockade by THZ531 elicited a non-genotoxic stress response that may drive the activation of P-TEFb. Giving credence to this premise, treatment of HCT116 cells with triptolide, which inhibits Pol II transcription by blocking the ATPase activity of the TFIIH subunit XPB (66), also led to the accumulation of p53 that occurred before increase in the levels of γ-H2AX-P (Figure 1E, right).

Because the immunoblotting approach might obscure more subtle increase in the levels of DNA damage in THZ531-treated cells, we asked whether inhibition of ataxia-telangiectasia mutated (ATM) protein kinase precludes the dissociation of CDK9 from HEXIM1. Namely, ATM is a chief orchestrator of the cellular DDR that gets activated by DSBs to regulate DNA repair, cell-cycle checkpoints and apoptosis by phosphorylation of multiple targets (67). Consistent with the γ-H2AX-P immunoblotting results, the dissociation of CDK9 remained unaffected upon co-treatment of HCT116 cells with THZ531 and a highly selective ATM inhibitor KU-60019 (68) for three hours. Later into the co-treatment, however, the inhibition of ATM enhanced, albeit slightly, the release of CDK9 from HEXIM1 by THZ531, which correlated with augmented levels of γ-H2AX-P (Figure 1F). Together, we conclude that inhibition of CDK12 activates P-TEFb by triggering its release from 7SK snRNP. While genotoxic stress does not appear to play a role early in this process, our results suggest that accumulating DNA damage in the course of prolonged inhibition of CDK12 contributes to a sustained and augmented activation of P-TEFb.

### P-TEFb promotes gene induction in CDK12-inhibited cells

Previous reports have established that inhibition of CDK12 leads to down-regulation as well as up-regulation of genes (12,16,23). However, in contrast to the down-regulated genes (9,10,22,23), the knowledge about the mechanisms that govern gene induction as well as the nature of the induced genes remains obscure. Considering our findings above, we hypothesized that activation of P-TEFb enables gene induction in CDK12-inhibited cells by stimulating the release of Pol II from promoter-proximal pause sites. To address this model, we utilized precision run-on sequencing (PRO-seq), which quantifies transcriptionally engaged Pol II complexes at nucleotide-resolution across the genome, including at promoter-proximal regions (69). We conducted these experiments using HCT116 *CDK12* ^as/as^ cell line, which allows for the selective inhibition of asCDK12 by 3-MB-PP1. To inhibit P-TEFb, we employed a sub-lethal, 10 nM dose of NVP-2, a highly selective inhibitor of P-TEFb that shows potent anti-cancer activity at non-toxic doses (70,71).

We treated HCT116 *CDK12* ^as/as^ cells with DMSO and 3-MB-PP1 or NVP-2 alone and in combination for 1 h and 4.5 h, and generated reproducible nucleotide-resolution maps of transcription with PRO-seq (Supplementary Figure S2A, B). We first quantified Pol II engagement at gene bodies, which are the sites of productive transcription elongation, to determine differentially transcribed genes (DTGs) using DESeq2 (*P*-adj ≤ 0.05; fold-change ≥ 1.25 for induced and ≤ 0.8 for down-regulated genes). Strikingly, we found that in comparison to DMSO treatment, increased engagement of Pol II dominated the altered transcriptional landscape at both durations of CDK12 inhibition. At 1 h, transcription of 25.3% of protein-coding genes (n = 16560) was changed, amongst which 3704 gained Pol II, while 478 were down-regulated (Figure 2A, left). At 4.5 h of CDK12 inhibition, this trend progressed in the favor of genes with gained Pol II. Out of 2164 DTGs, 2138 showed an increase in Pol II engagement, while only 26 genes showed a decrease (Figure 2A, left). Transcriptional induction triggered by the inhibition of CDK12 is exemplified at the long *DNMT1* gene, which demonstrates a wave of advancing Pol II that requires over an hour to reach the end of the gene (Supplementary Figure S2C). In contrast, the limited inhibition of P-TEFb alone reduced the engagement of Pol II along gene bodies as expected. While we found no DTGs at 1 h of P-TEFb inhibition, 3388 genes were down-regulated and 218 genes induced at 4.5 h of the treatment (Figure 2A, middle). Critically, inhibiting the activity of P-TEFb abrogated the induction of Pol II engagement elicited in CDK12-inhibited cells (Figure 2A, right). The block of Pol II entry into elongation occurred at 99.27% of CDK12 inhibition-induced genes already at 1 h of P-TEFb inhibition, shifting heavily the induced transcriptional landscape elicited by the inhibition of CDK12 towards down-regulation. This effect was even more striking at 4.5 h, when none of the 2138 genes induced in CDK12-inhibited cells showed a gain of Pol II. Intriguingly, we detected no down-regulated genes in the co-inhibited cells either, which is in sharp contrast to the effect observed in cells treated with the P-TEFb inhibitor alone. Highlighting that inhibition of CDK12 drastically changes the cellular transcriptional response to the limited P-TEFb inhibition (Figure 2A, middle and right), there is only a partial overlap between the P-TEFb-dependent genes of CDK12-inhibited cells (n = 2138) and control cells (n = 3388) at 4.5 h of the treatments (Supplementary Figure S2D). To this end, we conclude that P-TEFb promotes gene induction in CDK12-inhibited cells. Moreover, our results show that inhibition of CDK12 renders transcription of a new set of genes dependent on P-TEFb.

**Figure 2.**
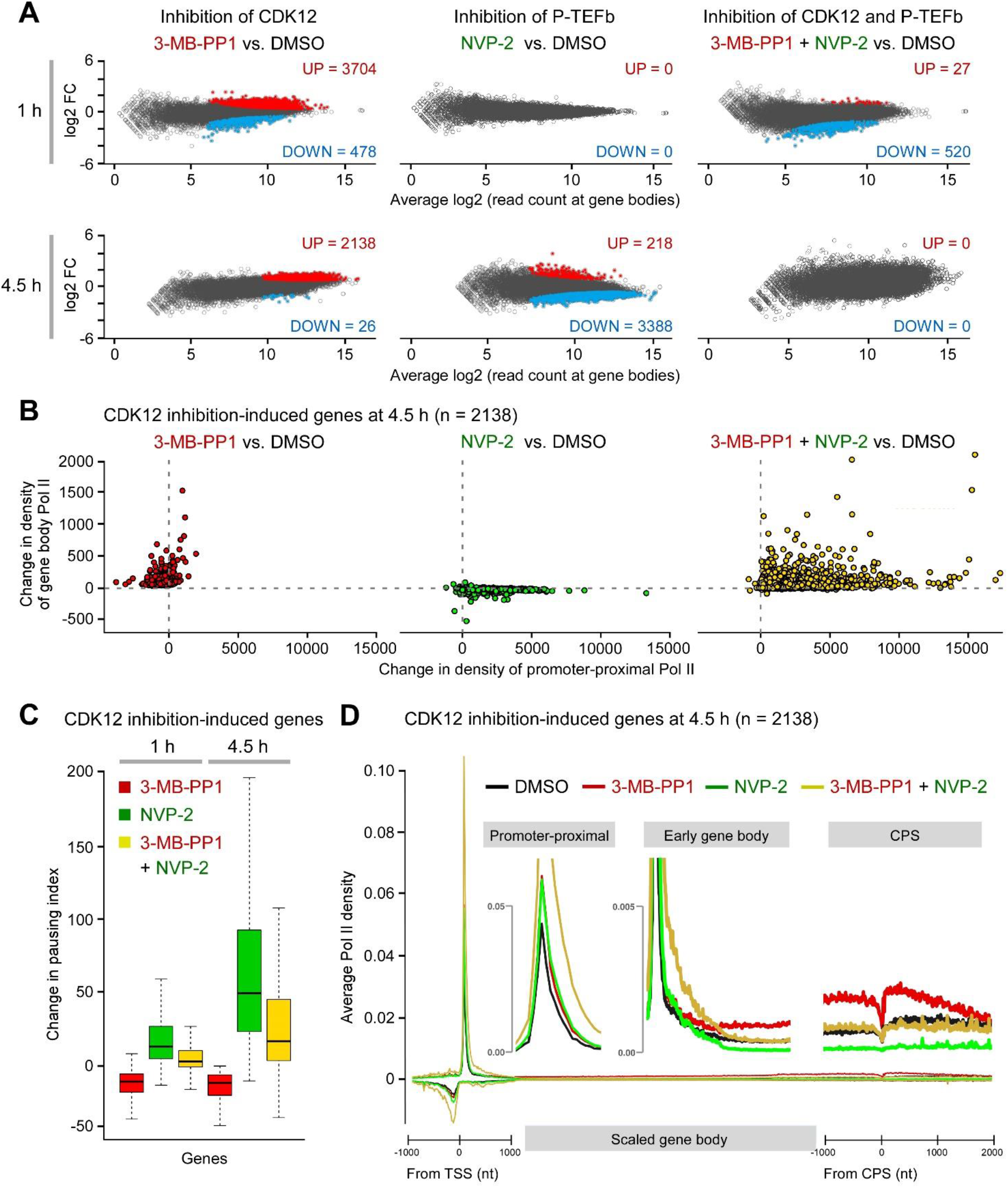
**Transcriptional induction in CDK12-inhibited cells occurs through P-TEFb-stimulated Pol II pause release.** (A) MA plot representation of differentially transcribed genes (n = 16,560; p-adj ≤ 0.05; FC ≥ 1.25 and ≤ 0.8) from PRO-seq experiments (n = 2) of HCT116 *CDK12* ^as/as^ cells treated with 3-MB-PP1 (5 μM) and NVP-2 (10 nM) as indicated for 1 h or 4.5 h compared to DMSO. Induced (UP), repressed (DOWN) and non-significantly regulated genes are depicted. FC, fold-change. (B) Change in the density of Pol II at promoter-proximal and gene body regions at CDK12 inhibition-induced genes at 4.5 h from PRO-seq experiments (n = 2) of (A). (C) Change in pausing index at CDK12 inhibition-induced genes at 1 h and 4.5 h from PRO-seq experiments (n = 2) of (A). Median pausing index for each group is indicated. (D) Average density of engaged Pol II at CDK12 inhibition-induced genes at 4.5 h from PRO-seq experiments (n = 2) of (A). The promoter-proximal region is measured as a linear scale from –1000 to +1000 nucleotide from the TSS, gene body scaled into 50 bins per gene, and end of the gene as a linear scale from –1000 to + 2000 nucleotide from the CPS. The insets show magnified promoter-proximal, early gene body and CPS gene regions. TSS, transcription start site. CPS, cleavage and polyadenylation site. nt, nucleotide.

### Inhibition of CDK12 induces transcriptional dependency on P-TEFb by stimulating Pol II pause release

To attain a mechanistic understanding for the induced transcription in CDK12 inhibited cells, we plotted the change in Pol II engagement at promoter-proximal and gene body regions of the induced genes at both durations of CDK12 and P-TEFb inhibition. This analysis revealed that increased Pol II engagement across the gene bodies in CDK12-inhibited cells was accompanied by decreased Pol II engagement at promoter-proximal regions at most genes (Figure 2B, left). These findings show that inhibition of CDK12 stimulates the escape of Pol II from the promoter-proximal pause, which stimulates transcription along gene bodies. Concomitantly, transcription initiation might also get increased at these genes as released Pol II from the pause has been shown to drive re-initiation of transcription (72). As expected, inhibition of P-TEFb alone increased Pol II engagement at pause sites, and blocked Pol II from entering productive elongation along the gene bodies (Figure 2B, middle). Critically, inhibition of P-TEFb countered the effects of CDK12 inhibition by retaining Pol II at the promoter-proximal region of virtually every induced gene (Figure 2B, right), demonstrating that CDK12 inhibition induced Pol II pause release via P-TEFb. The increased Pol II engagement at the promoter-proximal pause upon the double-inhibition was concomitant with the Pol II gain at gene bodies, suggesting an inefficient transition of Pol II into productive elongation. To quantify effects of both inhibitors on the efficiency of Pol II release from pausing into elongation, we next derived changes in pausing indices at both time points by dividing engagement of Pol II at promoter-proximal region with that across gene body. This approach revealed that inhibition of CDK12 decreased Pol II pausing index, while inhibition of P-TEFb opposed this effect (Figure 2C and Supplementary Figure S2E), highlighting again that P-TEFb drives stimulation of Pol II pause release at the induced genes upon the inhibition of CDK12.

Finally, we generated meta-gene profiles to track Pol II progression through the induced genes at 4.5 h of the treatments. Compared to the DMSO control, inhibition of P-TEFb increased the engagement of Pol II at promoter-proximal pause, and reduced transcription along the gene body below the DMSO control (Figure 2D), recapitulating the known phenotype of inhibited Pol II pause release. In contrast, inhibition of CDK12 stimulated efficient progression of Pol II from the promoter-proximal pause into productive elongation (Figure 2D, middle inset), demonstrating the opposing effects of CDK12 and P-TEFb inhibitors on transcription at these genes. Critically, the induced engagement of Pol II beyond the early gene body region in CDK12-inhibited cells was attenuated by the concurrent inhibition of P-TEFb (Figure 2D, middle inset). Of note, co-inhibition of both kinases elevated the engagement of Pol II at promoter-proximal region, suggesting that more Pol II complexes entered the pause, but were precluded from transitioning efficiently into gene bodies due to the block imposed by the inhibited P-TEFb. Indeed, after the early gene body region, the average density of engaged Pol II in co-inhibited cells fell below the one observed in CDK12-inhibited cells, equaling the level detected in DMSO-treated cells (Figure 2D, middle and right insets). Together, we conclude that by activating P-TEFb, inhibition of CDK12 stimulates gene transcription at the level of Pol II pause release, which in turn induces transcriptional dependency on P-TEFb.

### Inhibition of CDK12 induces gene expression of hallmark cancer signaling pathways

As CDK12 plays multiple roles in gene expression, which include promotion of elongation and processivity of Pol II (10-13,22,23), pre-mRNA splicing (15,16,19) and regulation of mRNA stability (16), the stimulation of Pol II pause release upon inhibition of CDK12 provides only an opportunity for genes to be induced. To define differentially expressed protein-coding genes (DEGs) upon selective inhibition of CDK12, we utilized gene expression fold-changes of nuclear RNA sequencing (RNA-seq) data from serum-synchronized HCT116 *CDK12* ^as/as^ cells treated for 4.5 h with DMSO or 3-MB-PP1 (23). Applying our criteria for differential gene expression (*P*-adj ≤ 0.001; log2 fold-change ≥ 1), we identified 12.32% DEGs (n = 12495). Of these, 304 genes were induced while 1235 were down-regulated (Figure 3A). Consistent with previous studies on transcriptional response to CDK12 inhibition and genotoxic stress (10,48,49,73), the induced genes were considerably shorter than the repressed genes (Supplementary Figure S3A).

**Figure 3.**
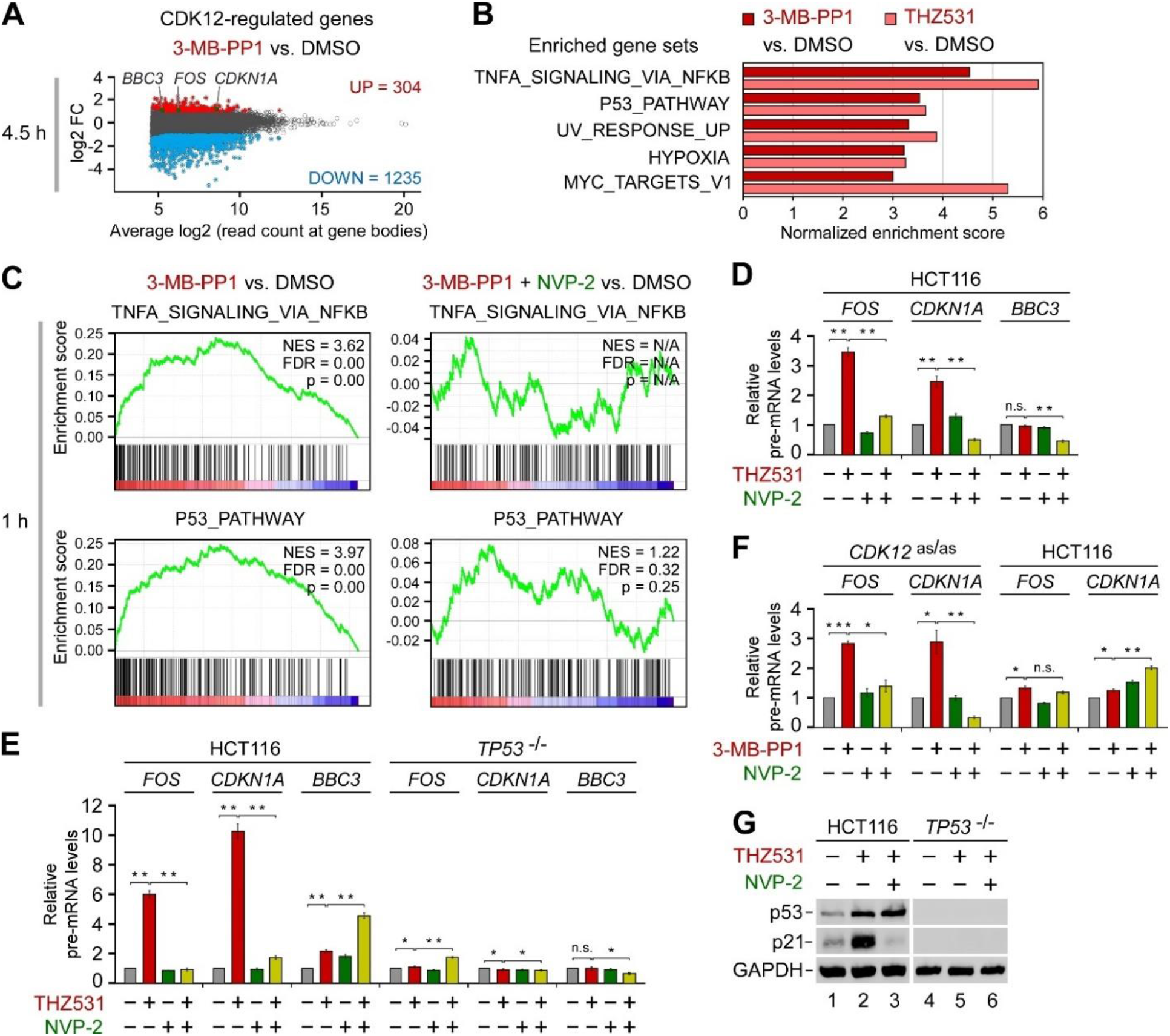
**Inhibition of CDK12 induces genes downstream of key cancer pathways via P-TEFb.** (A) MA plot representation of differentially expressed genes (n = 12,495; p-adj ≤ 0.001; Log2 FC ≥ 1) from nuclear RNA-seq experiments (n = 3) of HCT116 *CDK12* ^as/as^ cells that were synchronized by serum deprivation for 72 h prior to their culture in the serum-containing medium for 4.5 h in the presence of DMSO or 3-MB-PP1 (5 μM). Induced (UP), repressed (DOWN) and non-significantly regulated genes are depicted. *BBC3*, *FOS* and *CDKN1A* model genes are highlighted. FC, fold-change. (B) GSEA of gene expression changes for protein-coding genes of (A) and in A-375 cells treated with DMSO and THZ531 (500 nM) for 6 h. Top gene sets from the hallmark gene set collection that are associated with induced genes are shown (FDR q-val ≤0.001). (C) Enrichment plots of the top NF-κB and p53 pathway gene sets of (B) from the GSEA of transcription changes of protein-coding genes obtained from PRO-seq experiments (n = 2) of HCT116 *CDK12* ^as/as^ cells treated with DMSO, 3-MB-PP1 (5 μM) and NVP-2 (10 nM) as indicated for 1 h. NES, normalized enrichment score. FDR, false discovery rate. (D-F) HCT116 cell lines were treated with DMSO (-), THZ531 (400 nM), 3-MB-PP1 (5 μM), NVP-2 (20 nM) and KU-60019 (5 μM) alone and in combination as indicated for 3 h (D, F) or 12 h (E) prior to quantifying pre-mRNA levels of *FOS*, *CDKN1A* and *BBC3* with RT-qPCR. Results normalized to the levels of GAPDH mRNA and DMSO-treated cells are presented as the mean ± s.e.m. (n = 3). *, P < 0.05; **, P < 0.01; ***, P < 0.001, n.s., non-significant, determined by Student’s *t* test. (G) HCT116 cell lines were treated with DMSO (-), THZ531 (400 nM) and NVP-2 (20 nM) alone and in combination as indicated for 16 h prior to preparation of whole cell extracts and detection of the indicated proteins by Western blotting.

To gain an insight into the nature of the induced genes, we conducted gene set enrichment analysis (GSEA) of gene expression log2 fold-changes in the treated HCT116 *CDK12* ^as/as^ cells using the hallmark gene set collection of the Molecular Signatures Database (54,55). This analysis revealed that the top five gene sets enriched amongst the induced genes represented key oncogenic pathways, including NF-κB, hypoxia and MYC (Figure 3B and Supplementary Table S2A). In addition, the genes of the tumor suppressive p53 pathway and those induced in response to ultraviolet (UV) irradiation featured prominently, consistent with our immunoblotting results above and previous work that reported stimulation of p53 in cells with genetic or pharmacological perturbation of CDK12 (22,28,74). These findings were not specific to the CDK12-inhibited HCT116 *CDK12* ^as/as^ cells. Namely, we conducted GSEA of gene expression changes from multiple CDK12-inhibited cancer cell lines, including THZ531- treated melanoma A-375, Colo829 and neuroblastoma IMR-32 cell lines, and triple-negative breast cancer MDA-MB-231 cell line treated with CDK12 inhibitor SR-4835 (10,28,34), which revealed enrichment of the same gene sets (Figure 3B and Supplementary Table S2B-E). Likewise, GSEA of our HCT116 *CDK12* ^as/as^ PRO-seq data uncovered that the top NF-κB and p53 gene sets are enriched amongst the genes induced by the inhibition of CDK12 (Figure 3C and Supplementary Figure S3B, left). Importantly, blocking the activity of P-TEFb with NVP-2 either abrogated or significantly lowered the enrichment of the NF-κB gene set at 1 h or 4.5 h of CDK12 inhibition, respectively (Figure 3C and Supplementary Figure S3B, right). In contrast, while the addition of NVP-2 decreased the enrichment of the p53-dependent genes at 1 h, it failed to do so at 4.5 h into the treatment (Figure 3C and Supplementary Figure S3B, right). This result is in line with the reported ability of p53, of which levels accumulate during CDK12 inhibition (Figure 1E), to enable expression of a subset of its pro-apoptotic target genes under the limited inhibition of P-TEFb (42).

To validate these analyses, we used RT-qPCR to quantify pre-mRNA levels of the pro-survival *FOS* and *CDKN1A* as well as pro-apoptotic *BBC3* model genes. While both pro-survival genes, which encode the c-Fos subunit of the oncogenic transcription factor AP-1 and the CDK inhibitor p21, belong to the hallmark NF-κB, p53 and hypoxia pathway gene sets, *BBC3* is a classical p53 target gene, which encodes p53-upregulated modulator of apoptosis (PUMA) that belongs to BCL-2 protein family. We performed these experiments in the parental HCT116 cell line and in its isogenic *TP53* ^−/−^ derivative, in which *TP53* was inactivated by disrupting exon 2 of both alleles (75). In correspondence to our transcriptome analyses and the kinetics of CDK9 release from HEXIM1, treatment of HCT116 cells with 300 or 400 nM of THZ531 led to a time-dependent induction of *FOS* and *CDKN1A*, whereas 400 nM of THZ532 failed to do so (Supplementary Figure S3C). While the pro-survival genes showed considerable transcriptional induction early into the treatment, all three genes were induced at twelve hours of THZ531 exposure in a p53-dependent manner (Figure 3D, E). Importantly, the induction of *FOS* and *CDKN1A* was blunted by the addition of a sub-lethal dose of NVP-2 at three and twelve hours of the treatment (Figure 3D, E). We confirmed these results by selective inhibition of CDK12 with 3- MB-PP1 in HCT116 *CDK12* ^as/as^ cells, wherein both genes were induced in a P-TEFb-dependent manner (Figure 3F). In contrast, the induced pre-mRNA levels of *BBC3* were augmented by NVP-2 (Figure 3E), which is consistent with our earlier work that demonstrated differential response of p53- dependent genes, including *CDKN1A* and *BBC3*, to sub-lethal inhibition of P-TEFb (42). Finally, we conducted immunoblotting analysis using the parental and *TP53* ^−/−^ HCT116 cell lines to assess whether inhibition of CDK12 increases the expression of *CDKN1A* in a manner that requires P-TEFb and p53. In line with the transcript analyses, THZ531 treatment led to increased levels of p21 in a P-TEFb-and p53-dependent manner (Figure 3G). Together, we conclude that inhibition of CDK12 induces expression of genes downstream of important pathways in cancer, wherein P-TEFb plays a key role.

### Inhibition of CDK12 elevates the dependence of cancer cells on P-TEFb

Given our findings gathered thus far, we hypothesized that selective targeting of CDK12 renders cancer cells highly dependent on P-TEFb. As the hallmark p53 pathway gets activated in CDK12-inhibited cells, we further postulated that the status of p53 could play a key role in this process. We therefore utilized the parental and *TP53* ^−/−^ HCT116 cell lines and measured their response to selective inhibitors of CDK12 and P-TEFb.

We first employed fluorescence-based cytotoxicity assay, in which compromised membrane integrity of dead cells is proportional to fluorescence signal generated by the binding of an otherwise cell impermeable cyanine dye to DNA. We performed combinatorial titrations of THZ531 with NVP-2, where the highest dose of each inhibitor alone elicited maximal toxicity in HCT116 cells as established previously (42). We found that sub-lethal inhibition of CDK12 by THZ531 sensitized the parental cells to sub-lethal inhibition of P-TEFb by NVP-2 and iCDK9 (Figure 4A, C). We recapitulated this effect, when we replaced NVP-2 with iCDK9, which is another highly selective and ATP-competitive inhibitor of P-TEFb activity (76) (Supplementary Figure S4A). In contrast, the cytotoxic response to the co-targeting was largely diminished in the absence of p53 (Figure 4B, C and Supplementary Figure S4A), highlighting that activation of the p53-driven transcriptional program sensitizes CDK12-inhibited HCT116 cells to P-TEFb inhibitors. Next, we calculated Bliss synergy scores for all combinations using Synergyfinder (77). This approach identified the combination treatment of 200 nM of THZ531 with 10 nM of NVP-2 as the most synergistic in the parental HCT116 cells, reaching the Bliss score of 18 (Figure 3A, bottom). By comparison, the Bliss score of this co-treatment dropped by 6.9-fold in *TP53* ^−/−^ HCT116 cells (Figure 3B, bottom). In accordance with our p53-dependent cytotoxicity results and a key role of the kinase activity of ATM in the activation of p53 (78), inhibition of ATM by KU-60019 abrogated the dependence of CDK12-inhibited HCT116 cells on P-TEFb (Figure 4D). Importantly, in contrast to HCT116 cells, co-inhibition of CDK12 and P-TEFb was significantly less toxic for a non-transformed colorectal epithelial cell line CCD 841 CoN that encodes wild-type p53 (Supplementary Figure S4B), suggesting that cancer cells exhibit heightened dependence on CDK12 and P-TEFb. Confirming our results with THZ531, selective inhibition of CDK12 with 3-MB-PP1 in HCT116 *CDK12* ^as/as^ cells augmented cytotoxicity elicited by NVP-2 or iCDK9, whereas this effect was absent in the parental HCT116 cells (Figure 4E and Supplementary Figure S4C).

**Figure 4.**
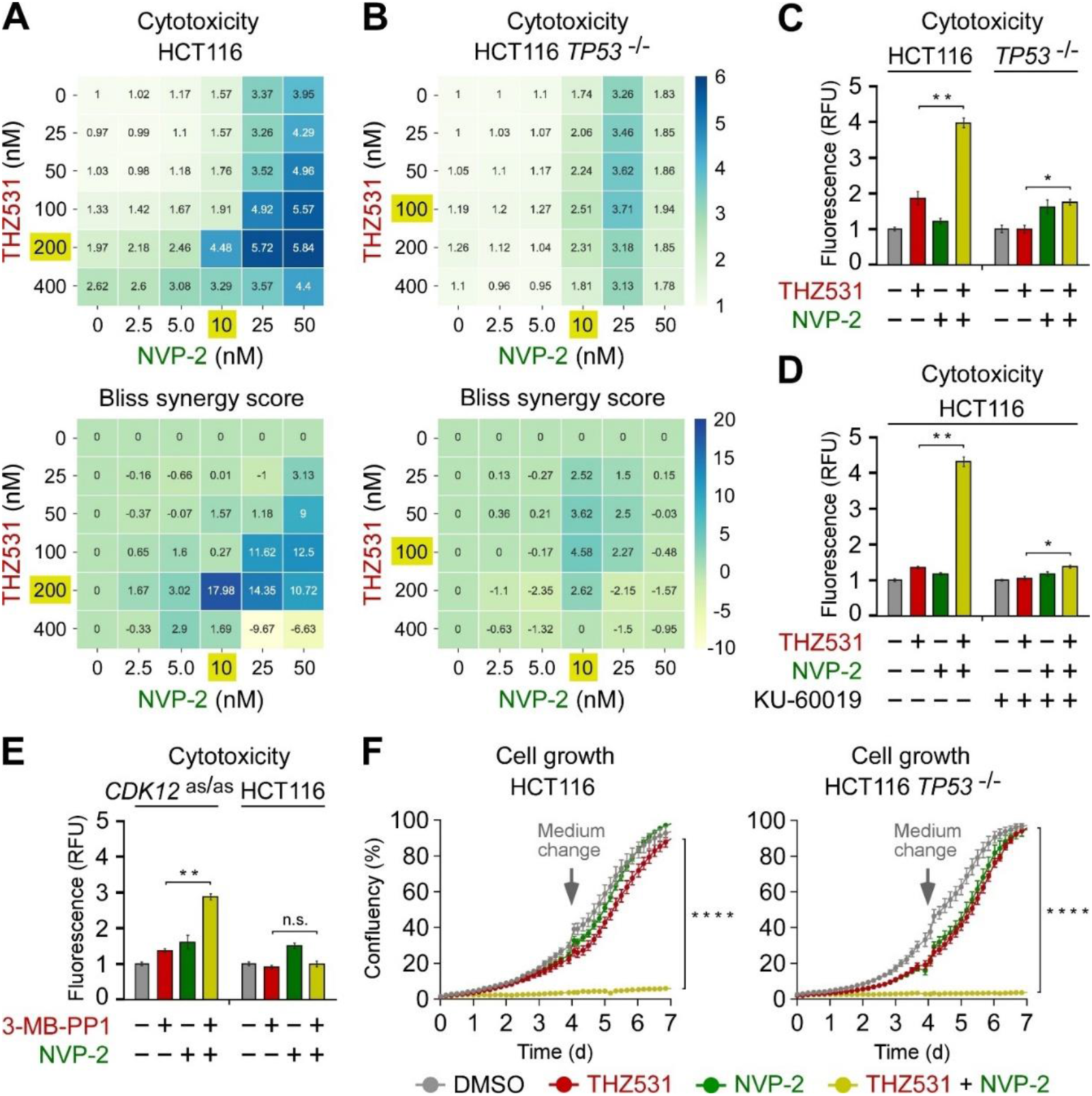
**Co-targeting of CDK12 and P-TEFb decreases viability of cancer cells.** (A,B) 6 × 6 cytotoxicity matrices with combinatorial titrations of THZ531 with NVP-2 at indicated doses to test for the synthetic lethality of compounds in HCT116 cell lines, depicting cytotoxicity (top) and synergy (bottom) of the combinations. Cytotoxicity values obtained at 48 hr of the treatments using CellTox Green assay were normalized to the DMSO control. Results represent the average of independent experiments (n = 3). Combinations with the highest Bliss synergy scores are highlighted (gold). (C-E) Cytotoxicity of HCT116 cell lines treated with DMSO (-), THZ531 (200 nM), 3-MB-PP1 (5 μM), NVP-2 (10 nM) and KU-60019 (5 μM) alone and in combination as indicated for 48 h measured using CellTox Green Cytotoxicity Assay. Results are presented as fluorescence values relative to the values of DMSO-treated cells and plotted as the mean ± s.e.m. (n = 3). *, P < 0.05; **, P < 0.01; n.s., non-significant, determined by Student’s *t* test. (F) Cell growth of HCT116 cell lines treated with DMSO, THZ531 (50 nM) and NVP-2 (1.25 nM) alone and in combination as indicated for seven days measured using live cell imaging. Results are presented as % confluency and plotted as the mean ± s.e.m. (n = 3). Arrows indicate the replenishment of medium and drugs at day 4. ****, P < 0.0001, determined by two-way ANOVA using THZ531 and THZ531 + NVP-2 data sets.

To probe the veracity of our cytotoxicity findings, we conducted the same experiments using melanoma A-375 and high-grade serous ovarian carcinoma Kuramochi cell lines, which differ in their status of p53. Whereas the former expresses functional p53 (79), the latter encodes a dominant negative p53, which contains the inactivating D281Y mutation located in its DNA-binding domain (80,81). Validating our HCT116 cell line results, co-inhibition of CDK12 and P-TEFb was highly toxic to p53-proficient A-375 cells but significantly less so to p53-deficient Kuramochi cells (Supplementary Figure S4D).

To provide complementary evidence for the synergistic lethality of CDK12 and P-TEFb co-targeting, we exposed the parental and *TP53* ^−/−^ HCT116 cell lines to sub-lethal doses of THZ531 and NVP-2 alone or in combination, and monitored cell growth during the seven day period using live cell imaging. While treatment of HCT116 cells with THZ531 and NVP-2 alone affected them similarly to the DMSO control, concurrent inhibition of CDK12 and P-TEFb resulted in a synergistic drop in cell growth (Figure 4F, left). Surprisingly, this effect was independent of p53 (Figure 4F, right), suggesting that an additional pathway elicited by CDK12 inhibition can also augment cellular dependency on P-TEFb. In parallel, we monitored metabolic activity of both HCT116 cell lines treated for up to seven days using a luminescence-based assay, in which the level of ATP quantified via luciferase reaction is proportional to the number of metabolically active cells and used as a measure of cell viability. Corroborating the cell growth results, inhibition of CDK12 by THZ531 and P-TEFb by NVP-2 or iCDK9 alone affected both cell lines minimally, while co-inhibition of the kinases decreased cell viability in a synergistic and p53- independent fashion (Supplementary Figure S4E, F). We confirmed these results using the asCDK12 system, in which co-administration of 3-MB-PP1 and NVP-2 or iCDK9 decreased viability of HCT116 *CDK12* ^as/as^ but not the parental cells (Supplementary Figure S4G, H). Consistent with our HCT116 cell line findings, co-inhibition of the kinases decreased viability of p53-deficient Kuramochi cells (Supplementary Figure S4I). Together, we conclude that selective inhibition of CDK12 renders HCT116 cells highly dependent on P-TEFb. Moreover, our results indicate that the status of p53 is a key factor influencing the fate of CDK12 and P-TEFb co-inhibited cancer cells. Whereas the combination treatment stimulates death of p53-proficient cells, it hampers proliferation of p53-deficient cells.

### Co-targeting of CDK12 and P-TEFb stimulates cancer cell death by p53-dependent apoptosis

We investigated further the significance of the hallmark p53 pathway for the elevated reliance of HCT116 cells on P-TEFb following selective inhibition of CDK12. Upon activation, the DNA-binding p53 orchestrates diverse tumor-suppressive transcriptional programs, amongst which cell-cycle arrest and apoptosis are best-defined (82). Hence, we conducted cell-cycle and apoptosis assays in the absence or presence of selective inhibitors of CDK12 and P-TEFb. Given the ability of p53 to drive cytotoxicity of CDK12 and P-TEFb co-targeted cells (Figure 4 and Supplementary Figure S4), we postulated that stimulation of p53-dependent apoptosis is the culprit. To quantify apoptosis, we utilized a luminescence-based assay that measures apoptosis-induced exposure of phosphatidylserine on the outer surface of cell membrane through time. The assay uses two Annexin V fusion proteins each containing a fragment of NanoBit Luciferase, of which complementation is achieved upon binding of Annexin V with phosphatidylserine (83). In concordance with the above cytotoxicity and viability data, THZ531 or NVP-2 treatment of HCT116 cells alone elicited a defect in the G1/S phase transition of the cell-cycle and low levels of apoptosis (Figure 5A, B). On the contrary, we detected synergistic apoptotic response of cells exposed to the combination treatment, which depended on p53 and was initiated at about twelve hours into the treatment (Figure 5B). Likewise, we observed the same phenotype when we targeted P-TEFb with iCDK9 in place of NVP-2 (Supplementary Figure S5A).

**Figure 5.**
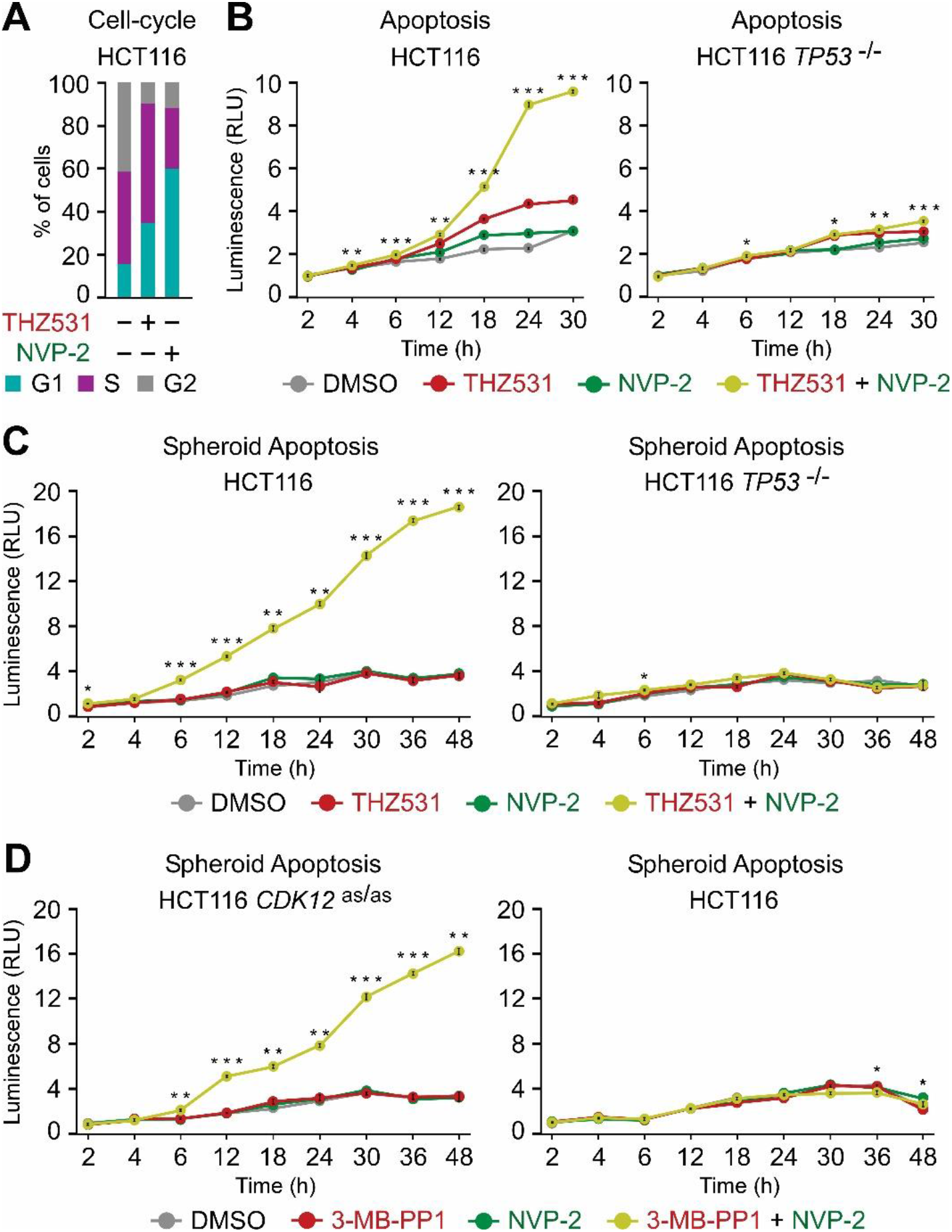
**Co-targeting of CDK12 and P-TEFb stimulates p53-dependent apoptosis of HCT116 cells.** (A) Quantification of cells (%) in the indicated cell-cycle phases based on flow cytometry profiles of propidium iodide-stained HCT116 cells that were synchronized by serum deprivation for 24 h prior to their culture in the serum-containing medium for 24 h in the presence of DMSO (-), THZ531 (50 nM) and NVP-2 (50 nM) as indicated. (B) Apoptosis of HCT116 cell lines treated with DMSO, THZ531 (100 nM) and NVP-2 (5 nM) alone and in combination as indicated. Results obtained at the time points indicated below the graphs using RealTime-Glo Annexin V Apoptosis and Necrosis Assay are presented as luminescence values relative to the values of DMSO-treated cells at 2 h and plotted as the mean ± s.e.m. (n = 3). *, P < 0.05; **, P < 0.01; ***, P < 0.001, determined by Student’s *t* test using THZ531 and THZ531 + NVP-2 data sets. (C,D) Apoptosis of HCT116 cell line spheroid cultures treated with DMSO, THZ531 (100 nM), 3-MB-PP1 (5 μM) and NVP-2 (10 nM) alone and in combination as indicated. Spheroids were formed for 48 h prior to the treatments. Results obtained at the time points indicated below the graphs using RealTime-Glo Annexin V Apoptosis and Necrosis Assay are presented as luminescence values relative to the values of DMSO-treated cells at 2 h and plotted as the mean ± s.e.m. (n = 3). *, P < 0.05; **, P < 0.01; ***, P < 0.001, determined by Student’s *t* test using THZ531 and THZ531 + NVP-2 (C), and 3- MB-PP1 and 3-MB-PP1 + NVP-2 (D) data sets.

To validate the apoptosis findings in a more physiological setting, we next assessed whether co-inhibition of CDK12 and P-TEFb stimulates apoptotic death of HCT116 spheroids. Namely, spheroid cultures have emerged as a viable pre-clinical platform for evaluating anti-cancer treatments (84). Unlike monolayer cultures, spheroids possess several *in vivo* hallmarks of tumor microenvironments such as spatial architecture of cells with different proliferation status, complex cell-cell contacts, oxygen and nutrient gradients, and rewired metabolism. As reported in our previous study (42), we established scaffold-free spheroids of about 400 μm in size before treating them for two days with DMSO, THZ531, NVP-2 or iCDK9 alone and in combination, during which we monitored kinetics of apoptosis utilizing the quantitative Annexin V assay. We observed that THZ531, NVP-2 and iCDK9 treatments in isolation elicited HCT116 spheroid apoptosis at vanishingly low levels that were indistinguishable from the DMSO control. In contrast, the co-targeting of CDK12 and P-TEFb elicited robust apoptosis in a highly synergistic manner (Figure 5C and Supplementary Figure S5B, left). As anticipated, the combination treatments failed to result in apoptosis of *TP53* ^−/−^ spheroids (Figure 5C and Supplementary Figure S5B, right). Likewise, 3-MB-PP1 treatment sensitized HCT116 *CDK12* ^as/as^ but not the parental HCT116 spheroids to P-TEFb inhibitors NVP-2 and iCDK9 (Figure 5D and Supplementary Figure S5C). Together, we conclude that inhibition of CDK12 primes cancer cells for p53-dependent apoptosis, which is triggered by co-inhibition of P-TEFb.

### Inhibition of CDK12 renders cancer cells dependent on the oncogenic NF-κB pathway

Because co-inhibition of CDK12 and P-TEFb decreased the growth of HCT116 cells independently of p53 (Figure 4), we reasoned that the targeting of CDK12 exposed novel vulnerabilities of cancer cells that could be exploited. To address this possibility, we turned our attention to the oncogenic pathways enriched within the genes induced by the inhibition of CDK12 (Figure 3B and Supplementary Table S2). We focused on the top-ranked NF-κB signaling pathway stimulated by TNF-α, of which enrichment diminishes upon inhibition of P-TEFb (Figure 3C and Supplementary Figure S3B). This canonical NF-κB pathway has emerged as a potential target in cancer. It promotes genesis and progression of cancer by facilitating proliferation, survival, inflammation, angiogenesis and invasion of tumor cells, and plays a role in the development of resistance to chemotherapy (85). Furthermore, the NF-κB pathway was the second most enriched pathway in tumors with hypertranscription, a recently revealed hallmark of aggressive human cancers (86). Importantly, NF-κB pathway can be stimulated from the nucleus by the DNA damage-induced activation of ATM (87). In both modes of the pathway activation (Figure 6A), key event is the signal-induced phosphorylation of the inhibitor of κB (IκB) by the IκB kinase β (IKKβ), which leads to IκB degradation. In turn, the cytoplasmic retention of NF-κB is alleviated, prompting stimulation of NF-κB-driven proliferative and anti-apoptotic gene transcription program.

**Figure 6.**
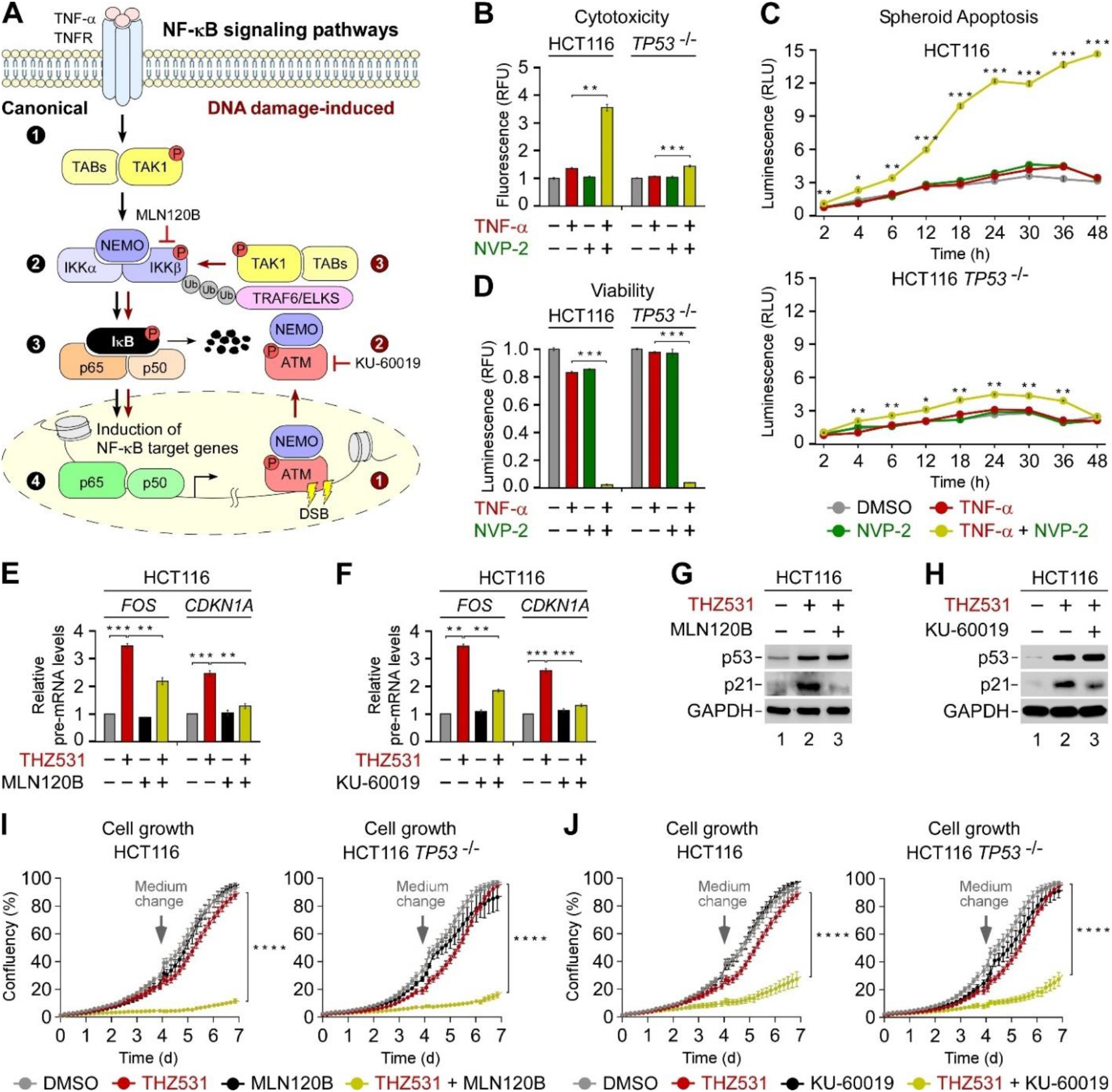
**Co-targeting of CDK12 and the NF-κB pathway decreases proliferation of HCT116 cells.** (A) Cartoon depicting canonical (left) and DNA damage-induced (right) NF-κB signaling pathway. In the canonical pathway, trimerization of TNFRSF1A by TNF-α leads to phosphorylation-induced activation of TAK1 (step 1), which in a complex with TABs phosphorylates and activates IKKβ of the IKK complex (step 2) that also includes a catalytic subunit IKKβ and a regulatory subunit NEMO/IKKγ. In turn, IKKβ induces the phosphorylation and degradation of IκBα (step 3), allowing NF-κB to translocate to the nucleus and stimulate proliferative and anti-apoptotic gene transcription program (step 4). In the DNA damage-induced pathway, induction of DSBs activates ATM and stimulates nuclear import of a fraction of NEMO (step 1), after which ATM in a complex with NEMO is exported from the nucleus (step 2) to stimulate TAK1-mediated phosphorylation and activation of IKKβ within signalosomes formed upon ATM-stimulated auto-polyubiquitylation of TRAF6 and polyubiquitylation of the IKK adaptor ELKS (step 3). Steps 3 and 4 of the canonical pathway ensue. Pharmacological targeting of IKKβ by MLN120B and ATM by KU-60019 is indicated. TNF-α, tumor necrosis factor α; TNFRSF1A, TNF receptor superfamily member 1A; TAK1, TGF-β-activated kinase 1; TABs, TAK1 binding proteins; NF-κB, nuclear factor kappa-light-chain-enhancer of activated B cells; IKKα, inhibitor of κB (IκB) kinase α; IKKβ, inhibitor of IκB kinase β; NEMO/IKKγ, NF-κB essential modulator; p65/p50, heterodimer of NF-κB; ATM, ataxia-telangiectasia mutated; TRAF6, tumor necrosis factor receptor– associated factor 6. (B) Cytotoxicity of HCT116 cell lines treated with DMSO, TNF-α (20 ng/ml) and NVP-2 (10 nM) alone and in combination as indicated for 48 h measured using CellTox Green Cytotoxicity Assay. Results are presented as fluorescence values relative to the values of DMSO-treated cells and plotted as the mean ± s.e.m. (n = 3). **, P < 0.01; ***, P < 0.001, determined by Student’s *t* test. (C) Apoptosis of HCT116 cell line spheroid cultures treated with DMSO, TNF-α (20 ng/ml) and NVP-2 (10 nM) alone and in combination as indicated. Spheroids were formed for 48 h prior to the treatments. Results obtained at the time points indicated below the graphs using RealTime-Glo Annexin V Apoptosis and Necrosis Assay are presented as luminescence values relative to the values of DMSO-treated cells at 2 h and plotted as the mean ± s.e.m. (n = 3). *, P < 0.05; **, P < 0.01; ***, P < 0.001, determined by Student’s *t* test using TNF-α and TNF-α + NVP-2 data sets. (D) Viability of HCT116 cell lines treated with DMSO, TNF-α (5 ng/ml) and NVP-2 (2.5 nM) alone and in combination as indicated for seven days measured using CellTiter-Glo 2.0 Cell Viability Assay. Results are presented as luminescence values relative to the values of DMSO-treated cells and plotted as the mean ± s.e.m. (n = 3). ***, P < 0.001, determined by Student’s *t* test. (E,F) HCT116 cell lines were treated with DMSO (-), THZ531 (400 nM), MLN120B (20 μM) and KU- 60019 (5 μM) alone and in combination as indicated for 3 h prior to quantifying pre-mRNA levels of *FOS* and *CDKN1A* with RT-qPCR. Results normalized to the levels of GAPDH mRNA and DMSO-treated cells are presented as the mean ± s.e.m. (n = 3). **, P < 0.01; ***, P < 0.001, determined by Student’s *t* test. (G,H) HCT116 cells were treated with DMSO (-), THZ531 (400 nM), NVP-2 (20 nM), MLN120B (20 μM) and KU-60019 (5 μM) alone and in combination as indicated for 16 h prior to preparation of whole cell extracts and detection of the indicated proteins by Western blotting. (I,J) Growth of HCT116 cell lines treated with DMSO, THZ531 (50 nM), MLN120B (2.5 μM) and KU- 60019 (50 nM) alone and in combination as indicated for seven days measured using live cell imaging. Results are presented as % confluency and plotted as the mean ± s.e.m. (n = 3). Arrows indicate the replenishment of medium and drugs at day 4. ****, P < 0.0001, determined by two-way ANOVA using THZ531 and THZ531 + MLN120B (I) and THZ531 and THZ531 + KU-60019 (J) data sets.

To determine contribution of this pathway to the lethality of CDK12 and P-TEFb co-targeting, we first asked whether its direct activation by TNF-α elevates the dependence of HCT116 cells on P-TEFb. Mirroring our findings in CDK12-inhibited cells, cytotoxicity assays revealed that TNF-α treatment sensitized HCT116 cells to P-TEFb inhibitors NVP-2 and iCDK9 in a p53-dependent manner (Figure 6B and Supplementary Figure S6A). Correspondingly, kinetic apoptosis measurements of the parental and *TP53* ^−/−^ HCT116 cell spheroids exposed to TNF-α and NVP-2 showed that unlike the individual treatments, the combination elicited apoptosis in a highly synergistic and p53-dependent fashion (Figure 6C). Furthermore, while the chronic treatment of both HCT116 cell lines with TNF-α and NVP-2 alone led to a modest decrease in cell viability as determined by ATP measurements, the combination treatment led to a synergistic drop that did not depend on p53 (Figure 6D). Thus, activation of the NF-κB pathway alone renders HCT116 cells sensitive to P-TEFb inhibition in a p53-independent manner.

We next addressed the significance of the NF-κB pathway for gene induction and cell fate in the context of CDK12-targeted HCT116 cells. To inhibit activation of the NF-κB pathway, we targeted it pharmacologically using IKKβ and ATM inhibitors MLN120B (88) and KU-60019, respectively. Indeed, addition of either of these compounds lowered the induction of *FOS* and *CDKN1A* upon treatment of HCT116 cells for three hours with THZ531 (Figure 6E, F). Likewise, the inhibition of IKKβ abrogated the induction of these genes in 3-MB-PP1-treated HCT116 *CDK12* ^as/as^ cells (Supplementary Figure S6B). Paralleling our findings with inhibition of P-TEFb at twelve hours into the treatments, the THZ531- induced levels of *FOS* and *CDKN1A* decreased in the presence of either MLN120B or KU-60019, while those of *BBC3* increased further (Supplementary Figure S6C, D). Correspondingly, the THZ531- induced levels of p21 were either blunted or significantly lowered by the inhibition of IKKβ or ATM, respectively (Figure 6G, H).

To decipher whether the fate of CDK12-inhibited HCT116 cells depends on activity of the oncogenic NF-κB pathway, we employed our battery of functional approaches. First, we treated the parental and *TP53* ^−/−^ HCT116 cell lines with sub-lethal doses of CDK12 and either IKKβ or ATM inhibitors alone or in combination, and monitored cell growth during the seven day period using live cell imaging. While treatment of HCT116 cells with THZ531 and MLN120B or KU-60019 alone affected them similarly to the DMSO control, the combination treatments ablated cell growth in a synergistic and p53-independent fashion (Figure 6I, J). Correspondingly, the inhibition of CDK12, IKKβ or ATM alone affected viability of the parental and *TP53* ^−/−^ HCT116 cells as determined by ATP measurements only slightly, but the CDK12-IKKβ and CDK12-ATM co-targeted cell lines exhibited profound loss of viability independently of p53 (Supplementary Figure S6E, F). Likewise, inhibition of CDK12 by 3-MB-PP1 rendered HCT116 *CDK12* ^as/as^ cells sensitive to the inhibition of NF-κB pathway by MLN120B (Supplementary Figure S6G). Finally, in contrast to the co-inhibition of CDK12 and P-TEFb, the CDK12-IKKβ and CDK12-ATM combination treatments were not toxic for HCT116 cells (Supplementary Figure S6H), demonstrating that attenuation of the NF-κB-dependent cell proliferation drives their potency. Together, we conclude that selective inhibition of CDK12 renders HCT116 cells highly dependent on the oncogenic NF-κB pathway, which occurs irrespectively of the status of p53.

### Inhibition of CDK12 or P-TEFb switches the fate of CDK7-inhibited cancer cells from cell-cycle arrest to apoptosis

That co-targeting of CDK12 and P-TEFb decreased viability of HCT116 cells prompted us to assess whether targeting of these critical transcriptional kinases in combination with CDK7 also yields the same phenotype. Apart from controlling gene transcription, cytoplasmic CDK7 is a master facilitator of cell-cycle progression by acting as a CAK for key cell-cycle CDKs, including CDK1, 2, and 4 (89). Hence, CDK7 represents an appealing target in cancer. Not surprisingly, selective inhibition of CDK7 in various cell lines or in the analog-sensitive HCT116 *CDK7* ^as/as^ setting results in cell-cycle arrest at the G1/S transition, in part by attenuation of the gene expression program driven by E2F (90,91).

As elimination of cancer cell by irreversible cell death is a desirable outcome of any effective cancer treatment (92,93), we examined whether the targeting of CDK12 or P-TEFb switches the fate of CDK7- inhibited cells from cell-cycle arrest to apoptosis. To inhibit CDK7, we employed YKL-5-124, a covalent inhibitor that targets cysteine 312 on CDK7 and shows over 100-fold greater selectivity for CDK7 over P-TEFb and CDK2 (91). In agreement with earlier reports (90,91), treatment of the parental and *TP53*^−/−^ HCT116 cells with YKL-5-124 was not cytotoxic. However, co-administration of THZ531 or NVP-2 sensitized HCT116 cells to CDK7 inhibition in a p53-dependent manner (Figure 7A, B). That p53 enables the death of CDK7 and CDK12 co-targeted cells is consistent with a previous report demonstrating that activation of the p53 transcriptional program sensitized cancer cells to CDK7 inhibitors (94). Importantly, the combination treatments failed to affect the non-transformed CCD 841 CoN cells (Supplementary Figure S7A, B). Corroborating these results, selective inhibition of CDK12 by 3-MB-PP1 in HCT116 *CDK12* ^as/as^ cells or P-TEFb by iCDK9 in HCT116 cells elicited toxicity in combination with CDK7 inhibition by YKL-5-124 (Supplementary Figure S7C, D).

**Figure 7.**
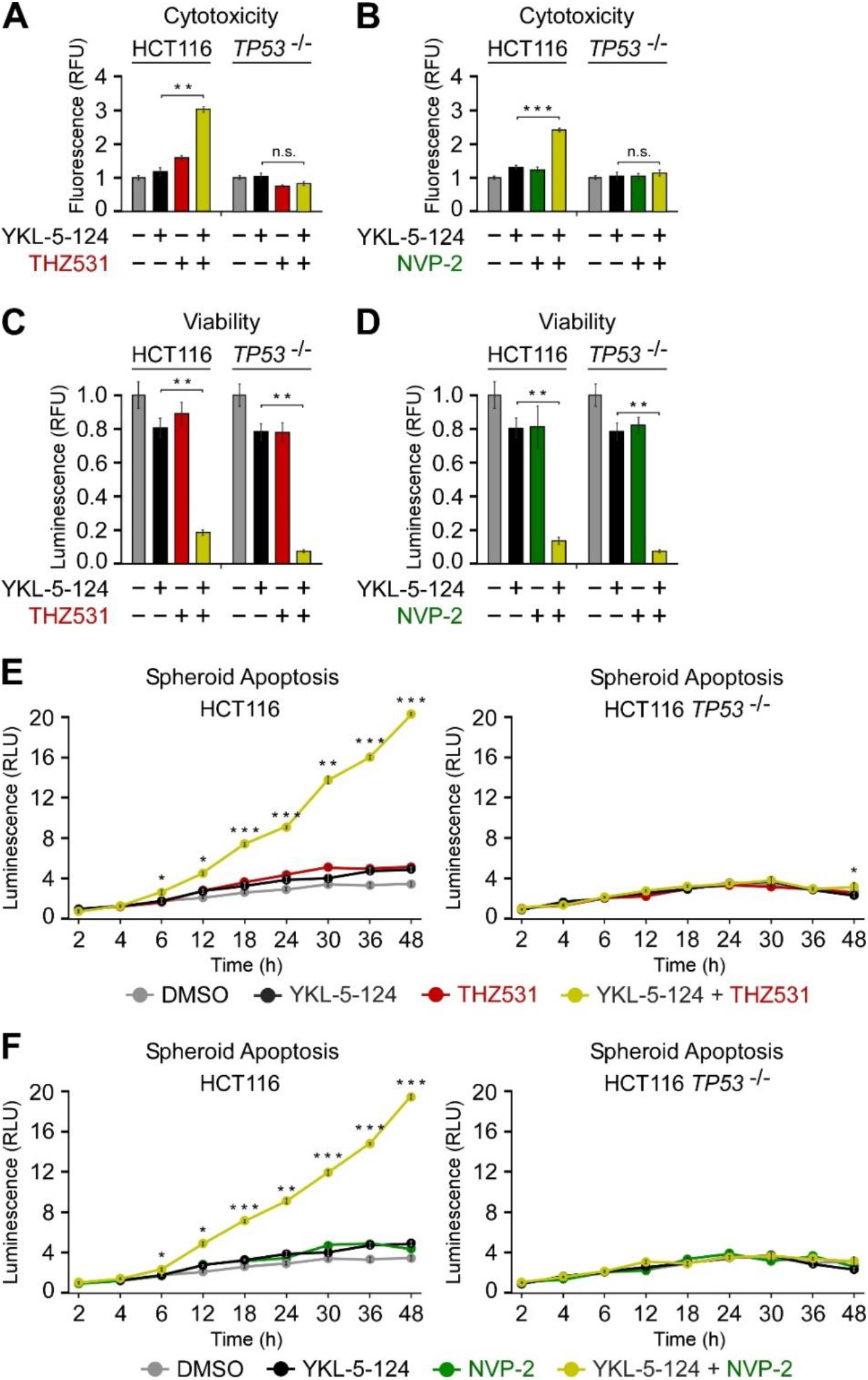
**Inhibition of CDK12 or P-TEFb switches the fate of CDK7-inhibited cancer cells from cell-cycle arrest to apoptosis.** (A,B) Cytotoxicity of the indicated HCT116 cell lines treated with DMSO, YKL-5-124 (100 nM), THZ531 (200 nM) and NVP-2 (10 nM) alone and in combination as indicated for 48 h measured using CellTox Green Cytotoxicity Assay. Results are presented as fluorescence values relative to the values of DMSO-treated cells and plotted as the mean ± s.e.m. (n = 3). *, P < 0.05; **, P < 0.01; ***, P < 0.001; n.s., non-significant, determined by Student’s *t* test. (C,D) Viability of the indicated HCT116 cell lines treated with DMSO, YKL-5-124 (5 nM), THZ531 (50 nM) and NVP-2 (2.5 nM) alone and in combination as indicated for seven days measured using CellTiter-Glo 2.0 Cell Viability Assay. Results are presented as luminescence values relative to the values of DMSO-treated cells and plotted as the mean ± s.e.m. (n = 3). **, P < 0.01, determined by Student’s *t* test. (E,F) Apoptosis of the indicated HCT116 cell line spheroid cultures treated with DMSO, YKL-5-124 (100 nM), THZ531 (150 nM) and NVP-2 (10 nM) alone and in combination as indicated. Spheroids were formed for 48 h prior to the treatments. Results obtained at the time points indicated below the graphs using RealTime-Glo Annexin V Apoptosis and Necrosis Assay are presented as luminescence values relative to the values of DMSO-treated cells at 2 h and plotted as the mean ± s.e.m. (n = 3). *, P < 0.05; **, P < 0.01; ***, P < 0.001, determined by Student’s *t* test using YKL-5-124 and YKL-5-124 + THZ531 (C), and YKL-5-124 and YKL-5-124 + NVP-2 (D) data sets.

To extend these findings, we conducted viability assays using the parental and *TP53* ^−/−^ HCT116 cells. Whereas treatment of HCT116 cells with the three tCDK inhibitors alone had a minimal effect on cell viability as determined by ATP measurements, co-administration of YKL-5-124 with THZ531 or NVP-2 led to a substantial and synergistic decrease of viability that did not require p53 (Figure 7C, D). Therefore, these cytotoxicity and viability results mirror our findings on the detrimental effect of CDK12 and P-TEFb co-targeting on cell fate.

Finally, we monitored apoptosis of the parental and *TP53* ^−/−^ HCT116 cell spheroids treated with the tCDK inhibitors in isolation or combination for 48 hours. While treatment of the spheroids with YKL-5- 124, THZ531 and NVP-2 alone stimulated apoptosis similarly to the DMSO control, co-administration of YKL-5-124 with THZ531 or NVP-2 elicited apoptosis in a highly synergistic and p53-dependent fashion (Figure 7E, F). Correspondingly, 3-MB-PP1 treatment sensitized HCT116 *CDK12* ^as/as^ but not the parental HCT116 spheroids to inhibition of CDK7 by YKL-5-124 (Supplementary Figure 7E). Likewise, co-targeting of CDK7 and P-TEFb by YKL-5-124 and iCDK9 led to a synergistic and p53- dependent stimulation of apoptosis (Supplementary Figure S7F). Collectively, these results confirm findings of the CDK-cyclin genetic co-dependencies of the DepMap project, where amongst cyclin genes, *CCNH* encoding the cyclin H of CDK7 is the top co-dependency of *CDK12*, whereas *CCNT1* encoding the cyclin T1 partner of CDK9 is the runner up co-dependency of CDK7 (20). Furthermore, they are in agreement with impressive anti-cancer activity of THZ1 in multiple in vivo models (64,95,96), wherein this analog of THZ531 co-inhibits CDK7 alongside CDK12/13 (63). Together, we conclude that selective inhibition of CDK12 or P-TEFb switches the fate of CDK7-inhibited HCT116 cells from cell-cycle arrest to apoptosis. However, further experimentation is needed to uncover the underlying mechanisms that drive the efficacy of the combinatorial treatments.

## DISCUSSION

An increasingly appreciated significance of tCDKs and deregulated transcription for cancer biology has put the prospects of therapeutic targeting of tCDKs into the spotlight. In this study, we selectively inhibited CDK12 by pharmacological and chemical genetic means in a well-established cell line model of colorectal cancer to uncover mechanistic and functional aspects of the induced transcriptome. We found that inhibition of CDK12 activated P-TEFb, which stimulated Pol II pause release and induced genes of prominent tumor suppressive and oncogenic signaling pathways exemplified by p53 and NF-κB, respectively. In turn, the CDK12-targeted cancer cells became hypersensitive to sub-lethal doses of selective inhibitors of P-TEFb. The limited inhibition of P-TEFb interfered with the gene induction in CDK12-targeted cells, and led to a synergistic elimination of these cells by stimulation of p53- dependent apoptosis and attenuation of NF-κB-dependent cell proliferation. Finally, the pairwise inhibition of CDK7 and either CDK12 or P-TEFb stimulated apoptosis of p53-proficient cancer cell spheroids. Therefore, we provide a rationale for combinatorial targeting of CDK12 with either P-TEFb or oncogenic signaling pathways in cancer.

Our study reveals a mechanism that governs the induced gene expression programs in CDK12- targeted cells. We demonstrate that inhibition of CDK12 provokes activation of P-TEFb through its release from the inhibitory 7SK snRNP reservoir, whereby most of the kinase leaves the complex within an hour of CDK12 inhibition. The kinetics of P-TEFb activation explains the progressive increase of the bulk of CTD Ser2 phosphorylation levels that was observed during six hours of inhibiting asCDK12 with 3-MB-PP1 (23). As demonstrated by our transcriptome-wide PRO-seq and model gene validation experiments, activated P-TEFb then stimulates the release of Pol II from promoter-proximal pausing, which culminates in transcriptional induction of short genes. As paused Pol II interferes with transcription initiation *in cis* (72), the P-TEFb-driven transition of Pol II into early elongation may enable subsequent bursts of Pol II initiation at induced genes, augmenting further their transcriptional output. Our findings are consistent with the prior work on THZ531-treated neuroblastoma cells, which showed a rapid and bidirectional accumulation of metabolically-labeled transcripts that originated from the start of transcription and extended well beyond the promoter-proximal pause (10). Likewise, they are reminiscent of the observed release of P-TEFb from 7SK snRNP following an acute degradation of the MED14 subunit of the Mediator complex, which augmented the rate of Pol II pause release genome-wide (97). It remains to be determined how exactly does the inhibition of CDK12 prompts the release of P-TEFb from 7SK snRNP. Based on our results, we propose that soon after the onset of CDK12 inhibition, global perturbation of transcription by Pol II elicits a non-genotoxic stress response that could drive the initial P-TEFb activation. Subsequently, prolonged inhibition of CDK12 results in increased load of DNA damage, which triggers cellular DDR, leading to sustained and augmented levels of P-TEFb activity.

Our findings clarify the nature and significance of the induced transcriptome in the wake of CDK12 inhibition. We show that the induced genes are enriched with genes stimulated by prominent cancer-related pathways, including p53, NF-κB, hypoxia and MYC. Supporting these findings, GSEA of hallmark pathways against the DepMap dataset of ranked genetic co-dependencies of *CDK12* across the genome revealed that the effect of *CDK12* knockout on cancer cell growth correlates positively with the oncogenic NF-κB and hypoxia pathways and negatively with the tumor suppressive p53 pathway (20). Furthermore, CDK12 is required for survival and transformation of MYC-overexpressing cells (24,98). By focusing on the p53 and NF-kB pathways, the chief elements of the cellular DDR, we reveal that gene induction in CDK12-inhibited cells is functionally important and a prerequisite for the increased sensitivity of cancer cells to P-TEFb inhibitors. In principle, decreased expression of CDK12- dependent genes could also contribute directly to the elevated dependence of CDK12-inhibited cells on P-TEFb. However, our recent work and findings in this study demonstrate that in the case of p53 and NF-κB pathways, gene induction is the culprit. Namely, we disentangled induced from repressed genes in CDK12-inhibited cells by selectively activating p53- and NF-κB-driven transcriptional programs using the non-genotoxic MDM2 inhibitor Nutlin-3a and TNF-α, respectively, which by itself renders the treated cells highly sensitive to P-TEFb inhibitors (42). Nevertheless, in the context of CDK12-targeted cells, the attenuated gene transcription could enable gene induction indirectly through the stress responses proposed above, which may prompt stimulation of P-TEFb and signal-responsive genes. Finally, a large body of work has established that Pol II promoter-proximal pausing is prevalent on signal-responsive genes, whereby transcriptional activators recruit P-TEFb to facilitate synchronous release of paused Pol II into productive elongation (99). Intriguingly, all TFs that operate at the endpoint of the top enriched pathways in CDK12-targeted cells, namely NF-κB, HIF1A, MYC and p53, have been reported to stimulate Pol II pause release through P-TEFb (100-103). Taken together, our findings are consistent with a model in which inhibition of CDK12 elicits cell stress-mediated activation of signaling pathways and P-TEFb, which leads to the capturing of this key Pol II pause release kinase by pathway-specific TFs on chromatin for coordinate gene activation. Accordingly, we envision that this sequence of events underpin the elevated dependence of CDK12-inhibited cells on P-TEFb.

Our work offers fresh avenues for combinatorial targeting of CDK12 in cancer. Thus far, down-regulation of CDK12-dependent HR genes is being explored as an approach to eliminate cancer cells, most notably in combination with PARP1 inhibitors or DNA-damaging chemotherapeutics (28,29). However, our results call for therapeutic exploitation of novel vulnerabilities and resistance mechanisms that arise in response to gene induction in CDK12-inhibited cancer cells. First, the targeting of CDK12 reactivates p53 and thus primes the cells for undergoing cell death by apoptosis, which we show can be triggered upon co-inhibition of P-TEFb or CDK7. These results resonate with reported studies demonstrating that activation of the p53 transcriptional program sensitizes cancer cells to selective inhibitors of P-TEFb or CDK7 (42,94). Second, our findings indicate that a host of oncogenic signaling pathways get induced in CDK12-inhibited cells, cautioning against the therapeutic targeting of CDK12 alone. However, by interrogating the DNA damage-induced NF-κB pathway, we provide a proof-of-concept for the combinatorial targeting of CDK12 with oncogenic signaling. We demonstrate that the stimulation of the NF-κB pathway alone by TNF-α triggers cell death in combination with P-TEFb inhibitors. Critically, we reveal that the pharmacological targeting of CDK12 in combination with either the inhibitors against P-TEFb and CDK7 or inhibitors of IKKβ and ATM of the DNA damage-induced NF-κB pathway leads to a synergistic drop of cancer cell proliferation irrespective of p53. These findings are highly relevant from a therapeutic standpoint as up to a half of cancer cases carry loss-of-function mutations in both alleles of *TP53*. Moreover, as activation of NF-κB has been linked to the resistance of ovarian cancer cells to PARP1 inhibitors (104), targeting this pathway could improve the outcome of the CDK12-PARP1 combination therapy. Finally, we show that the activity of p53 influences the seemingly paradoxical impact of ATM blockade on the fate of tCDK-inhibited cells. On the one hand, inhibition of ATM prevents the death of p53-proficient cells co-treated with CDK12 and P-TEFb inhibitors, which we attribute to the requirement of ATM for the activation of p53 (78). On the other hand, while blocking ATM is not lethal for CDK12-inhibited cells, it diminishes their proliferation independently of p53 by countering the oncogenic NF-κB pathway. Therefore, these results provide further credence to the fact that the functionality of *ATM* and *TP53*, two commonly mutated tumor suppressor genes, is a key determinant of the clinical response to genotoxic chemotherapies. Namely, while cancer cells with functional p53 display chemoresistance upon inhibition of ATM or its downstream target Chk2, inhibition of ATM sensitizes p53-deficient cancer cells to chemotherapy (105). Hence, we propose that the combined status of *ATM* and *TP53* is likely to dictate the efficacy of the CDK12-targeted combination therapies. Conversely, CDK12-deficient tumors could be particularly susceptible to the targeting of P-TEFb, CDK7 and the DNA damage-induced ATM-Chk2 pathway.

Finally, our work provides new insights into the tumor suppressive role of CDK12 in the genesis of cancers with biallelic *CDK12* inactivation such as in high-grade serous ovarian carcinoma (30-32). On the one hand, disruption of CDK12-dependent regulatory networks elicits genomic disarray with the characteristic tandem DNA duplications. On the other hand, our results suggest that *CDK12* loss of heterozygosity engenders the incipient cancer cells with augmented oncogenic signaling that drive key hallmarks of cancer, which in the case of NF-kB signaling include sustained proliferative signaling, avoidance of programmed cell death, tumor-promoting inflammation and therapy resistance (85). Therefore, future exploration of processes driven by perturbation of CDK12 shall reveal the full potential of therapeutic targeting of this crucial Pol II elongation kinase in cancer.

## DATA AVAILABILITY

All data associated with this study are presented in the main manuscript or the Supplementary Data. Materials that support the findings of this study are available from the corresponding author upon request. PRO-seq raw data and normalized bigwig files have been deposited to Gene Expression Omnibus (GSE236803).

## ACKNOWLEDGEMENTS

We thank Joaquin M. Espinosa for sharing parental and HCT116 *TP53* ^-/-^ cell lines; Dalibor Blazek for sharing HCT116 *CDK12* ^as/as^ cell line; Malgorzata Krajewska and Rani E. George for conducting the target engagement assay of Figure 1C; Qiang Zhou for sharing i-CDK9; Nathanael S. Gray for sharing YLK-5-124; HiLife Biomedicum Flow Cytometry Unit at the University of Helsinki for flow cytometry analysis; Monika Mačáková and all the authors for critical reading of the manuscript.

## AUTHOR CONTRIBUTIONS

M.B. conceptualized the study and was assisted by Z.W. in experimental design. M.B. and A.V. supervised the study. S.V.H. and A.V. conducted PRO-seq experiments with the assistance of Z.W., and analyzed the PRO-seq data. Z.W. performed and analyzed the data of all other experiments except the target engagement assay of Figure 1C. H.M.H. analyzed the live cell imaging data. C.C.F performed gene expression analyses of RNA-seq data and GSEA analyses of RNA-seq, PRO-seq and TT-seq data. M.B. wrote the manuscript with the assistance from A.V. and Z.W.

## Supplementary Data

**Contents:**

1. Supplementary Figures S1-S7

2. Supplementary Tables S1-S2

**Supplementary Figure S1.**
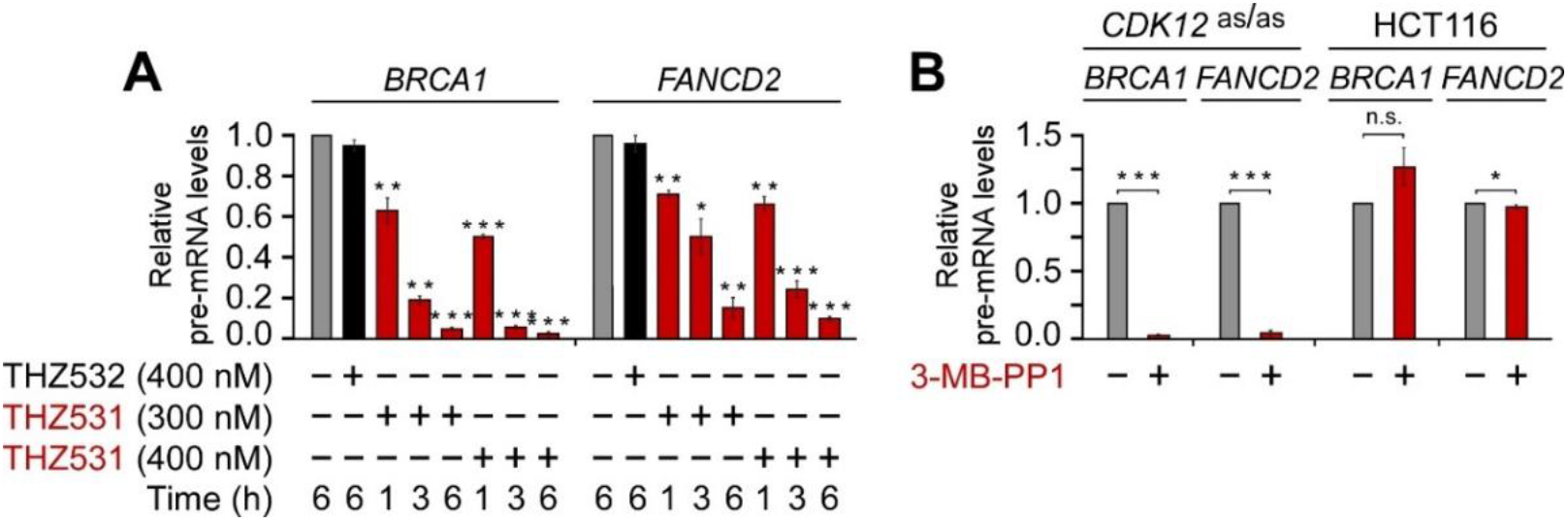
**Inhibition of CDK12 triggers the release of P-TEFb from the inhibitory 7SK snRNP complex.** (A) HCT116 cells were treated with DMSO (-), THZ532 and two different doses of THZ531 as indicated for the indicated duration prior to quantifying pre-mRNA levels of *BRCA1* and *FANCD2* with RT-qPCR. Results normalized to the levels of GAPDH mRNA and DMSO-treated cells are presented as the mean ± s.e.m. (n = 3). *, P < 0.05; **, P < 0.01; ***, P < 0.001, determined by Student’s *t* test. (B) HCT116 cell lines were treated with DMSO (-) and 3-MB-PP1 (5 μM) as indicated for 3 h prior to quantifying pre-mRNA levels of *BRCA1* and *FANCD2* with RT-qPCR. Results normalized to the levels of GAPDH mRNA and DMSO-treated cells are presented as the mean ± s.e.m. (n = 3). *, P < 0.05; ***, P < 0.001; n.s., non-significant, determined by Student’s *t* test.

**Supplementary Figure S2.**
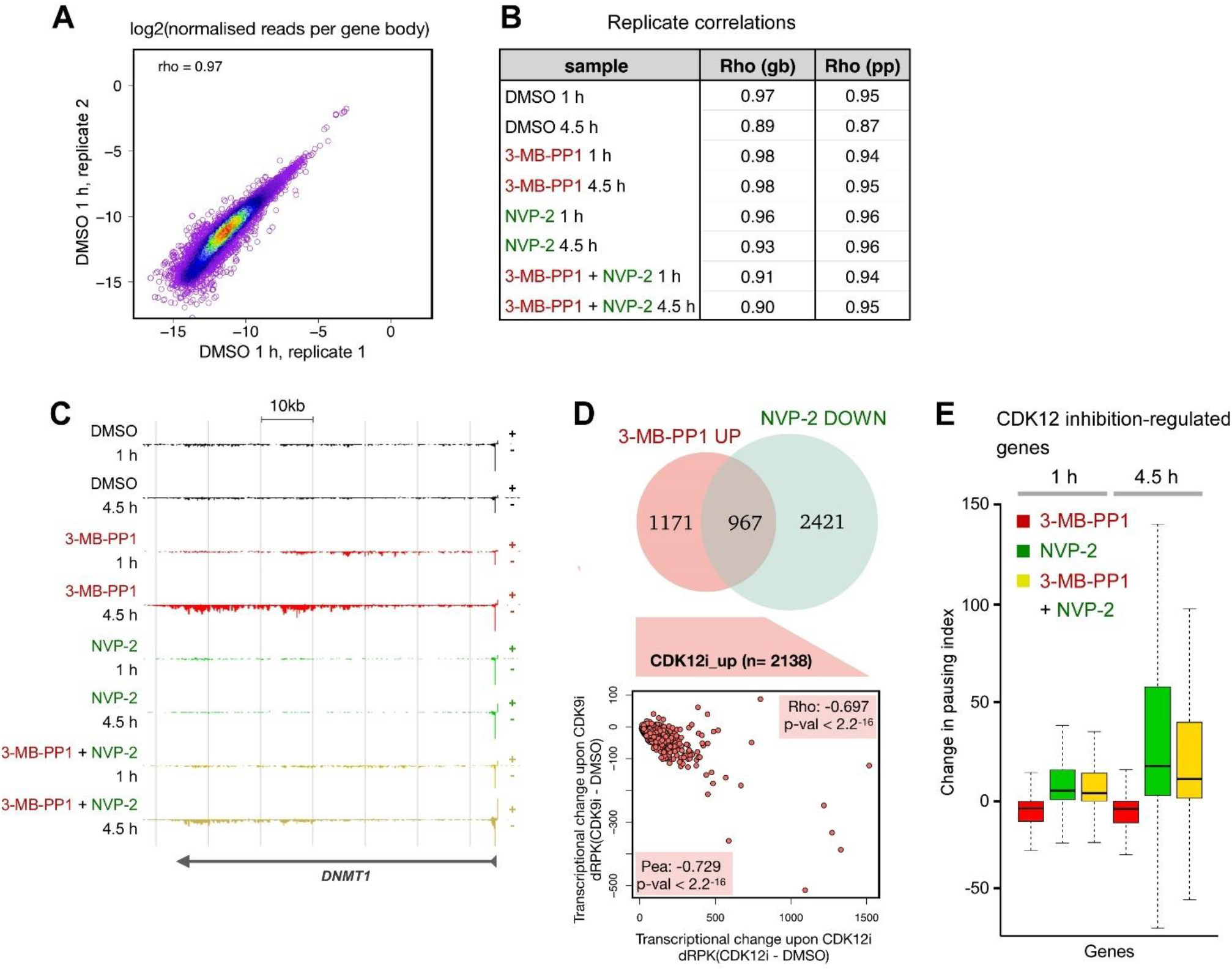
**Transcriptional induction in CDK12-inhibited cells occurs by P-TEFb-stimulated Pol II pause release.** (A) Representative correlation plot comparing PRO-seq replicates (n = 2) of HCT116 *CDK12* ^as/as^ cells treated with DMSO for 1 h. The x-and y-axes show log2 count of normalized PRO-seq reads across the gene bodies from +500 nt from the TSS to -500 nt from the TES. (B) Correlations between the PRO-seq replicates (n = 2) of HCT116 *CDK12* ^as/as^ cells treated with DMSO, 3-MB-PP1 (5 μM) and NVP-2 (10 nM) alone and in combination as indicated for 1 h and 4.5 h across promoter-proximal region (pp; -250 nt to +250 nt from the TSS) and gene body (gb) as defined in (A). Rho indicates Spearman’s rank correlation. (C) Density of engaged Pol II across *DNMT1* gene at 1 h or 4.5 h from PRO-seq replicates (n = 2) of HCT116 *CDK12* ^as/as^ cells treated with DMSO, 3-MB-PP1 (5 μM) and NVP-2 (10 nM) alone and in combination as indicated. Please note that the wave of transcriptional induction upon inhibition of CDK12 has proceeded 40 kb at 1 h, and through the entire gene at 4.5 h of the treatment. (D) Venn diagram (top) and correlation plot (bottom) comparisons between CDK12-induced and P-TEFb-repressed genes at 4.5 h of the PRO-seq replicated (n = 2) of HCT116 *CDK12* ^as/as^ cells treated with 3-MB-PP1 (5 μM) and NVP-2 (10 nM), respectively. Rho and Pea indicate Spearman’s and Pearson’s rank correlations, respectively. (E) Change in pausing index at CDK12 inhibition-regulated genes derived from PRO-seq replicates (n = 2) of HCT116 *CDK12* ^as/as^ cells treated with 3-MB-PP1 (5 μM) and NVP-2 (10 nM) alone and in combination as indicated for 1 h or 4.5 h compared to DMSO. Median pausing index for each group is indicated.

**Supplementary Figure S3.**
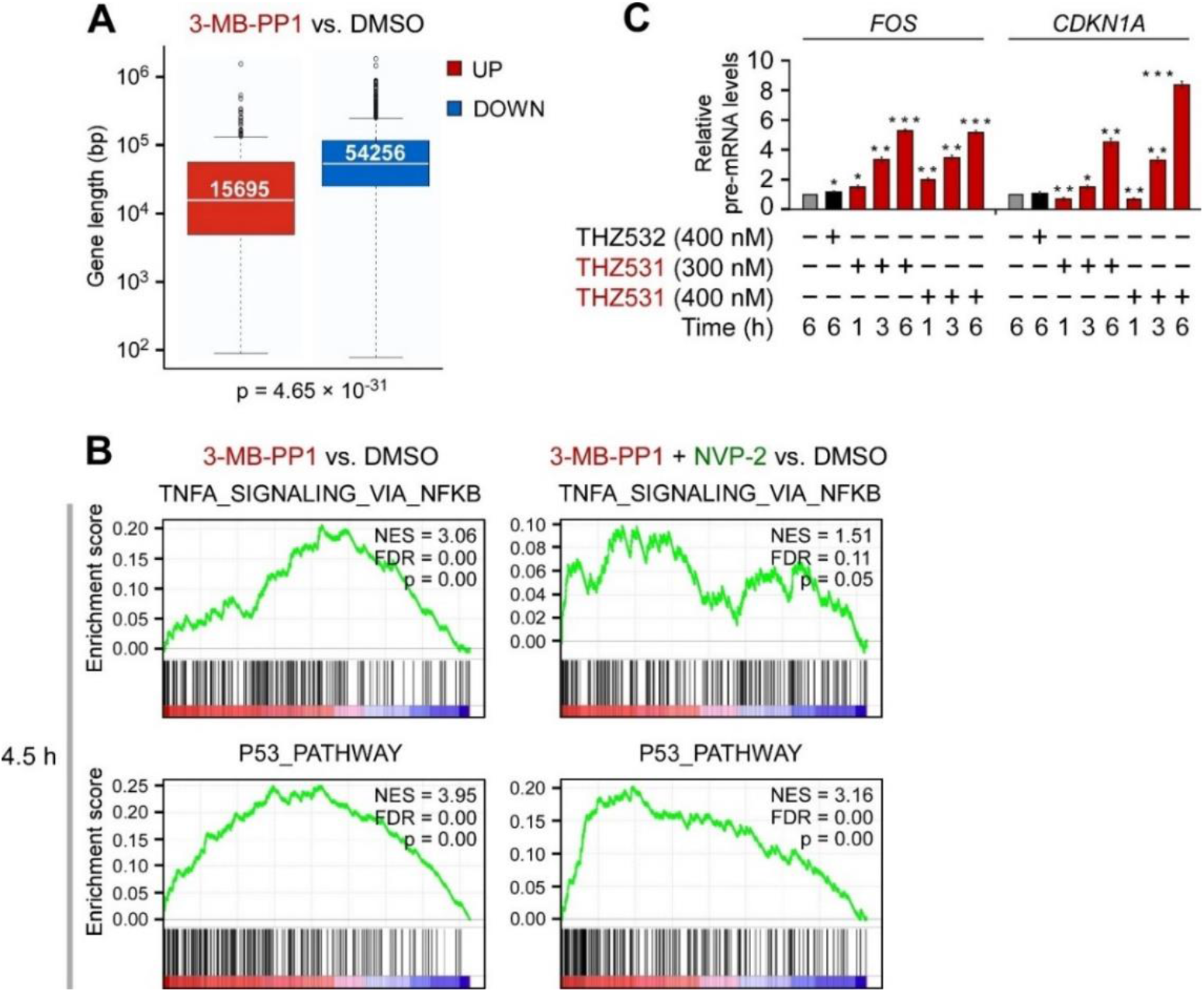
**Inhibition of CDK12 induces gene expression downstream of key cancer pathways.** (A) Boxplot indicating distribution of gene lengths for the induced (UP) and down-regulated (DOWN) protein-coding genes from RNA-seq experiments (n = 3) of serum-synchronized HCT116 *CDK12* ^as/as^ cells treated for 4.5 h with DMSO or 3-MB-PP1 (5 μM). Median gene length for each group and p-value are indicated. bp, base pair. (B) Enrichment plots of the top NF-κB and p53 pathway gene sets of Figure 3B from the GSEA of transcription changes of protein-coding genes obtained from PRO-seq experiments (n = 2) of HCT116 *CDK12* ^as/as^ cells treated with DMSO, 3-MB-PP1 (5 μM) and NVP-2 (10 nM) as indicated for 4.5 h. NES, normalized enrichment score; positive values indicate enrichment among induced genes, negative values enrichment among down-regulated genes. FDR, false discovery rate. (C) HCT116 cells were treated with DMSO, THZ532 and two different doses of THZ531 as indicated for the indicated duration prior to quantifying pre-mRNA levels of *FOS* and *CDKN1A* with RT-qPCR. Results normalized to the levels of GAPDH mRNA and DMSO-treated cells are presented as the mean ± s.e.m. (n = 3). *, P < 0.05; **, P < 0.01; ***, P < 0.001, determined by Student’s *t* test.

**Supplementary Figure S4.**
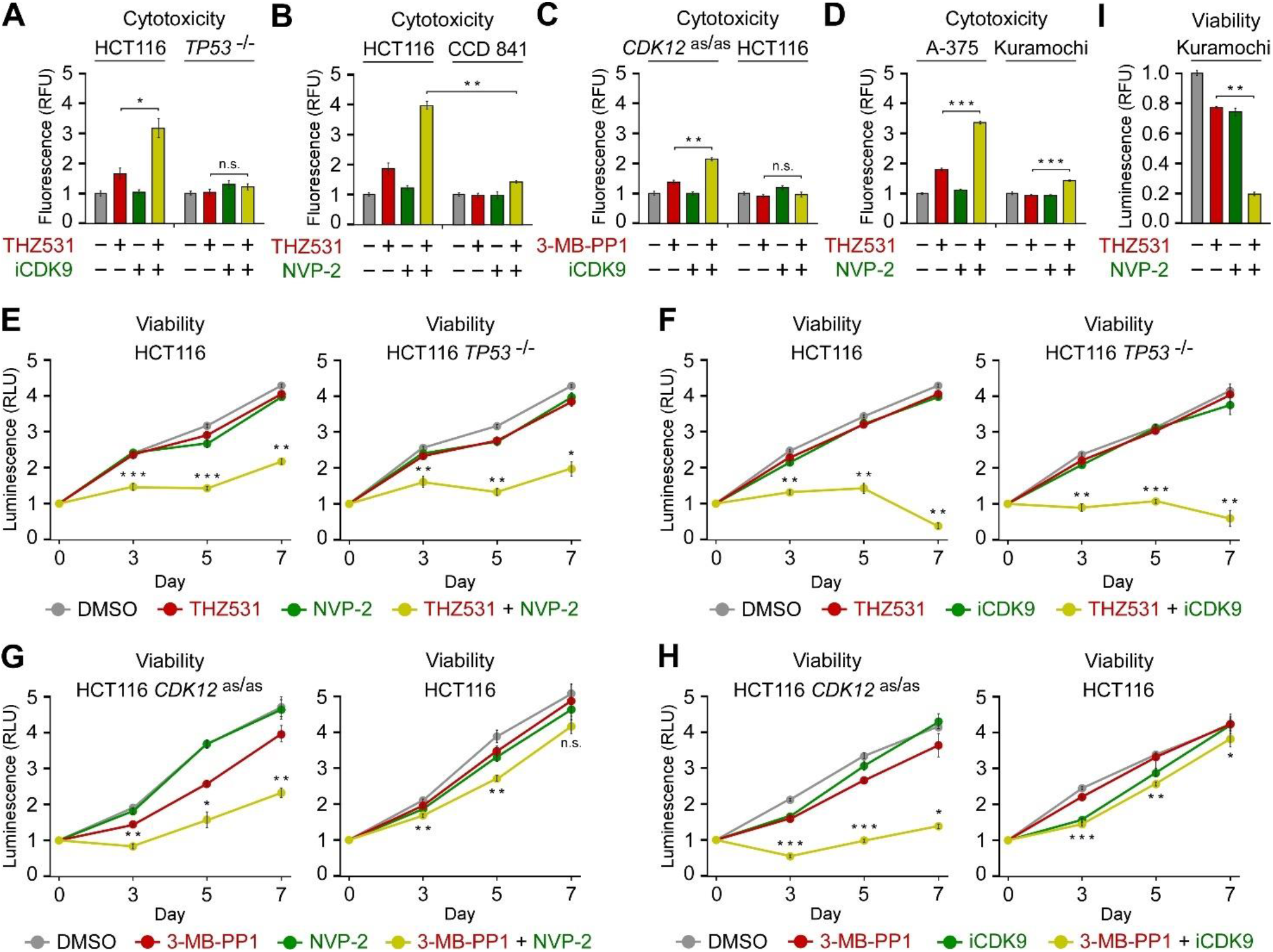
**Co-targeting of CDK12 and P-TEFb decreases viability of cancer cells.** (A-D) Cytotoxicity of the indicated cell lines treated with DMSO (-), THZ531 (200 nM), 3-MB-PP1 (5 μM) and iCDK9 (25 nM) alone and in combination as indicated for 48 h measured using CellTox Green Cytotoxicity Assay. Results are presented as fluorescence values relative to the values of DMSO-treated cells and plotted as the mean ± s.e.m. (n = 3). *, P < 0.05; **, P < 0.01; ***, P < 0.001; n.s., non-significant, determined by Student’s *t* test. (E,F) Viability of HCT116 cell lines treated with DMSO, THZ531 (50 nM), NVP-2 (2.5 nM) and iCDK9 (25 nM) alone and in combination as indicated. Results obtained at the time points indicated below the graphs using CellTiter-Glo 2.0 Cell Viability Assay are presented as luminescence values relative to the values at Day 0 and plotted as the mean ± s.e.m. (n = 3). *, P < 0.05; **, P < 0.01; ***, P < 0.001, determined by Student’s *t* test using THZ531 and THZ531 + NVP-2 (E), and THZ531 and THZ531 + iCDK9 (F) data sets. (G,H) Viability of HCT116 cell lines treated with DMSO, 3-MB-PP1 (2.5 μM), NVP-2 (1.25 nM) and iCDK9 (5 nM) alone and in combination as indicated. Results obtained at the time points indicated below the graphs using CellTiter-Glo 2.0 Cell Viability Assay are presented as luminescence values relative to the values at Day 0 and plotted as the mean ± s.e.m. (n = 3). *, P < 0.05; **, P < 0.01, ***, P < 0.001, determined by Student’s *t* test using 3-MB-PP1 and 3-MB-PP1 + NVP-2 (G), and 3-MB-PP1 and 3-MB-PP1 + iCDK9 (H) data sets. (I) Viability of Kuramochi cells treated with DMSO, THZ531 (50 nM) and NVP-2 (2.5 nM) alone and in combination as indicated. Results obtained at the time points indicated below the graphs using CellTiter-Glo 2.0 Cell Viability Assay are presented as luminescence values relative to the values at Day 0 and plotted as the mean ± s.e.m. (n = 3). **, P < 0.01, determined by Student’s *t* test using THZ531 and THZ531 + NVP-2 data sets.

**Supplementary Figure S5.**
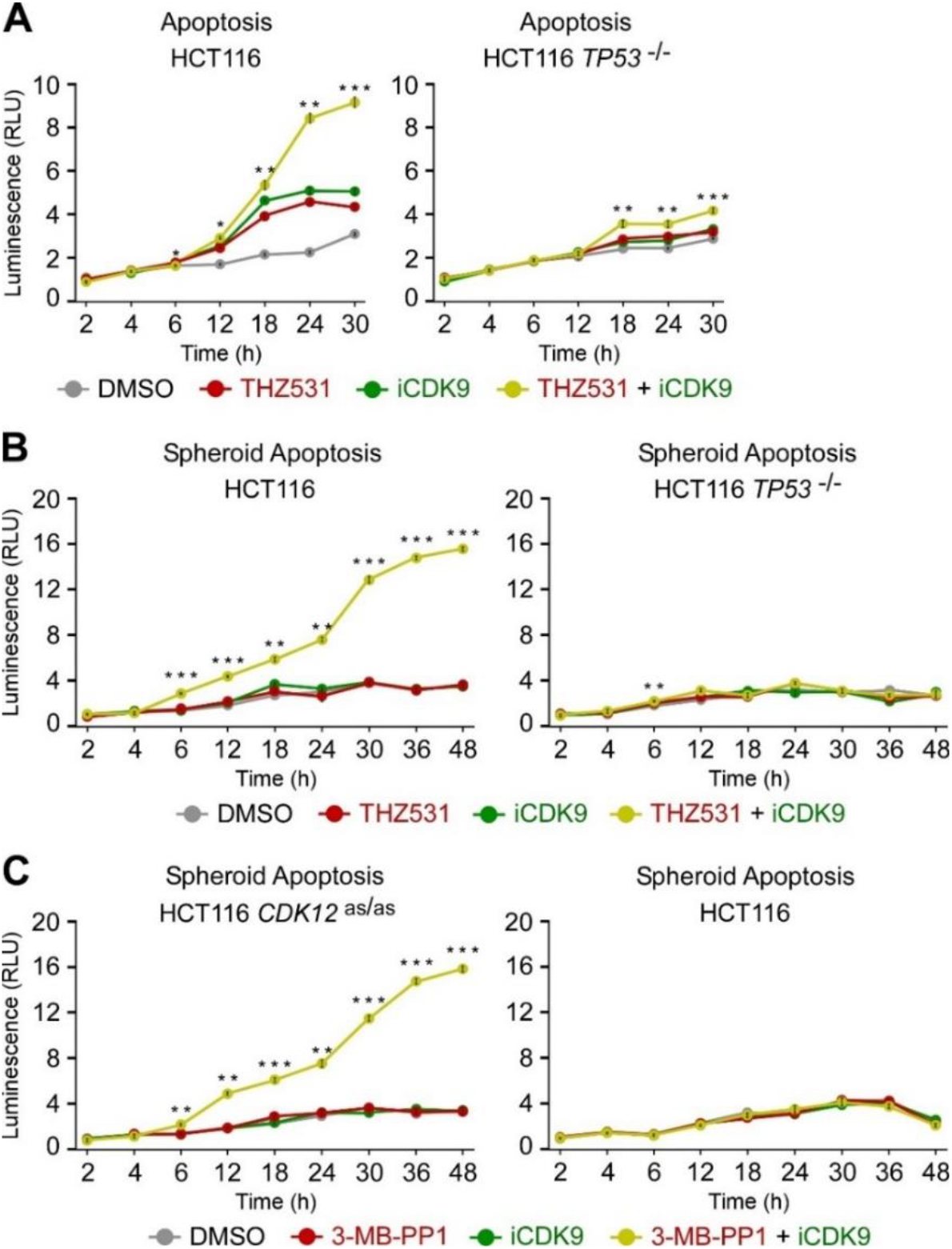
**Co-targeting of CDK12 and P-TEFb stimulates apoptosis of HCT116 cells.** (A) Apoptosis of HCT116 cell lines treated with DMSO, THZ531 (100 nM) and iCDK9 (25 nM) alone and in combination as indicated. Results obtained at the time points indicated below the graphs using RealTime-Glo Annexin V Apoptosis and Necrosis Assay are presented as luminescence values relative to the values of DMSO-treated cells at 2 h and plotted as the mean ± s.e.m. (n = 3). *, P < 0.05; **, P < 0.01; ***, P < 0.001, determined by Student’s *t* test using THZ531 and THZ531 + iCDK9 data sets. (B,C) Apoptosis of HCT116 cell line spheroid cultures treated with DMSO, THZ531 (100 nM), 3-MB-PP1 (5 μM) and iCDK9 (25 nM) alone and in combination as indicated. Spheroids were formed for 48 h prior to the treatments. Results obtained at the time points indicated below the graphs using RealTime-Glo Annexin V Apoptosis and Necrosis Assay are presented as luminescence values relative to the values of DMSO-treated cells at 2 h and plotted as the mean ± s.e.m. (n = 3). **, P < 0.01; ***, P < 0.001, determined by Student’s *t* test using THZ531 and THZ531 + iCDK9 (B), and 3-MB-PP1 and 3-MB-PP1 + iCDK9 (C) data sets.

**Supplementary Figure S6.**
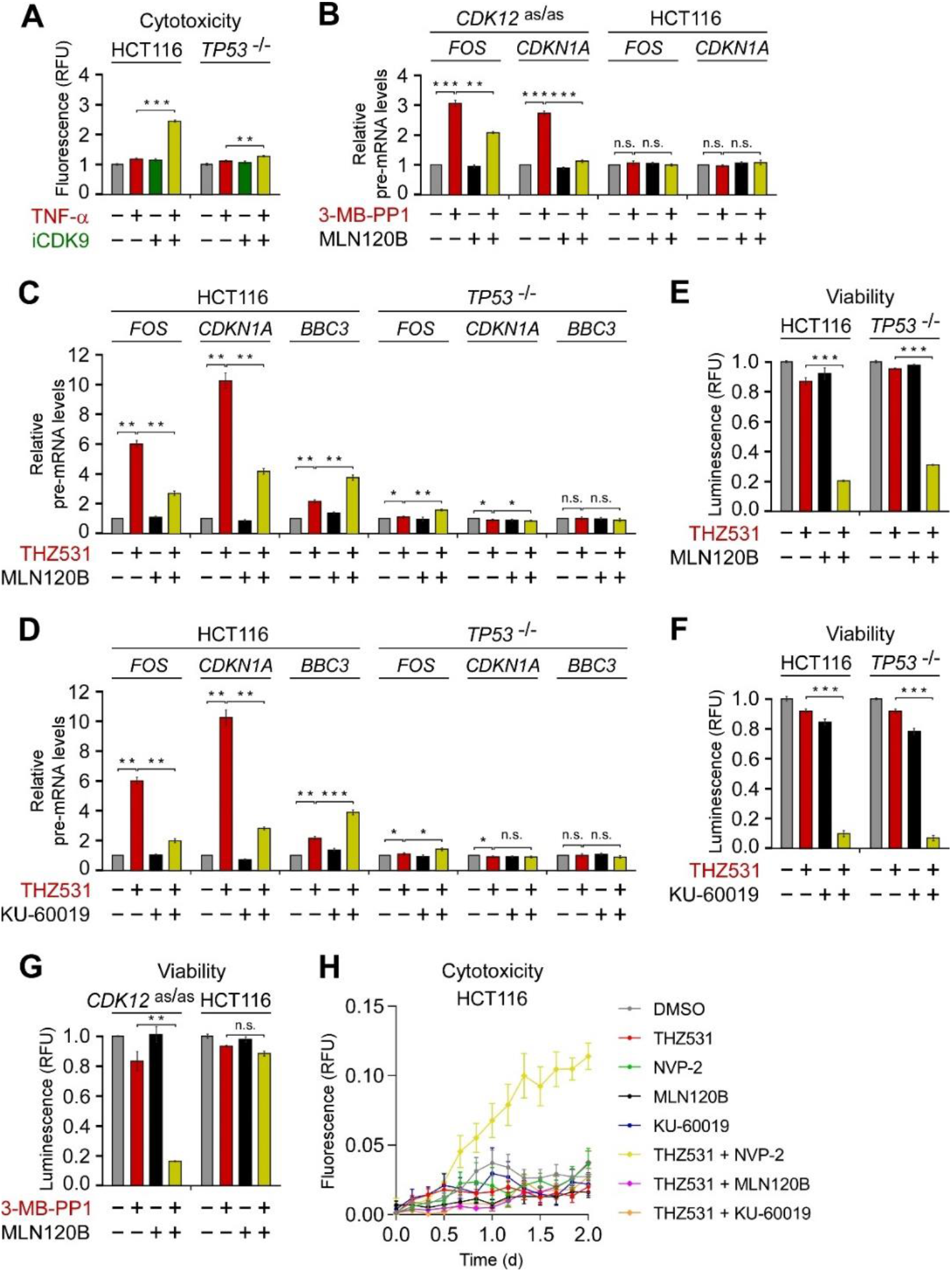
**Inhibition of CDK12 renders cancer cells dependent on the NF-κB pathway.** (A) Cytotoxicity of HCT116 cell lines treated with DMSO (-), TNF-α (20 ng/ml) and iCDK9 (25 nM) alone and in combination as indicated for 48 h measured using CellTox Green Cytotoxicity Assay. Results are presented as fluorescence values relative to the values of DMSO-treated cells and plotted as the mean ± s.e.m. (n = 3). **, P < 0.01; ***, P < 0.001, determined by Student’s *t* test. (B-D) HCT116 cell lines were treated with DMSO (-), THZ531 (400 nM), 3-MB-PP1 (5 μM), MLN120B (20 μM) and KU-60019 (5 μM) alone and in combination as indicated for 3 h (B) or 12 h (C, D) prior to quantifying pre-mRNA levels of *FOS*, *CDKN1A* and *BBC3* with RT-qPCR. Results normalized to the levels of GAPDH mRNA and DMSO-treated cells are presented as the mean ± s.e.m. (n = 3). *, P < 0.05; **, P < 0.01; ***, P < 0.001, n.s., non-significant, determined by Student’s t test. (E-G) Viability of HCT116 cell lines treated with DMSO, THZ531 (50 nM), 3-MB-PP1 (2.5 μM), MLN120B (2.5 μM in E; 5 μM in F) and KU-60019 (50 nM) alone and in combination as indicated for seven days measured using CellTiter-Glo 2.0 Cell Viability Assay. Results are presented as luminescence values relative to the values of DMSO-treated cells and plotted as the mean ± s.e.m. (n = 3). **, P < 0.01; ***, P < 0.001; n.s., non-significant, determined by Student’s t test. (H) Cytotoxicity of HCT116 cell lines treated with DMSO, THZ531 (50 nM), NVP-2 (1.25 nM), MLN120B (2.5 μM) and KU-60019 (50 nM) alone and in combination as indicated for 48 h measured using CellTox Green Cytotoxicity Assay. Results are presented as fluorescence values and plotted as the mean ± s.e.m. (n = 3).

**Supplementary Figure S7.**
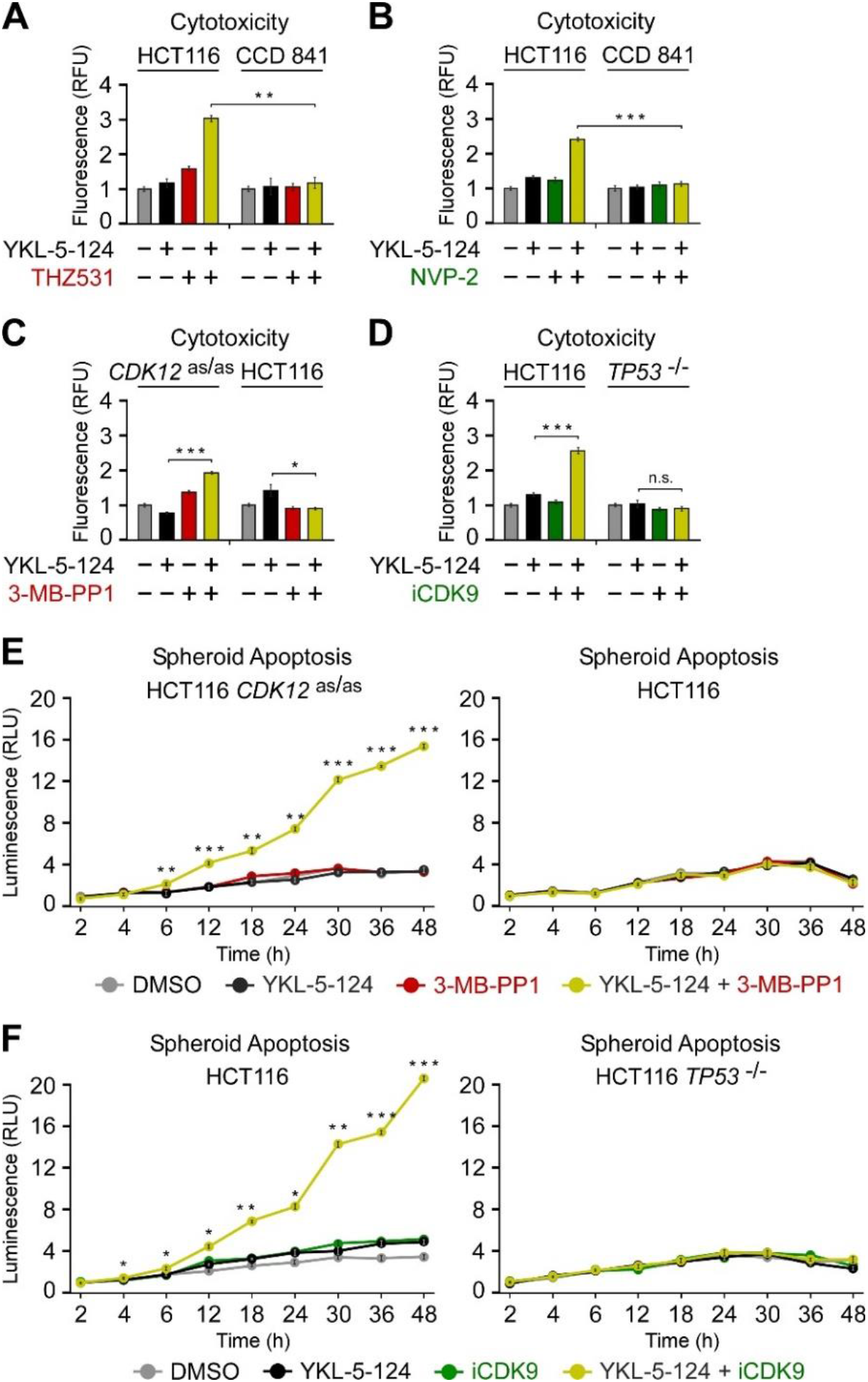
**Co-targeting of CDK7 and either CDK12 or P-TEFb stimulates apoptosis of HCT116 cells.** (A-D) Cytotoxicity of the indicated cell lines treated with DMSO (-), YKL-5-124 (100 nM), THZ531 (200 nM), 3- MB-PP1 (5 μM), NVP-2 (10 nM) and iCDK9 (25 nM) alone and in combination as indicated for 48 h measured using CellTox Green Cytotoxicity Assay. Results are presented as fluorescence values relative to the values of DMSO-treated cells and plotted as the mean ± s.e.m. (n = 3). **, P < 0.01; ***, P < 0.001; n.s., non-significant, determined by Student’s *t* test. (E,F) Apoptosis of HCT116 cell line spheroid cultures treated with DMSO, YKL-5-124 (100 nM), 3-MB-PP1 (5 μM) and iCDK9 (25 nM) alone and in combination as indicated. Spheroids were formed for 48 h prior to the treatments. Results obtained at the time points indicated below the graphs using RealTime-Glo Annexin V Apoptosis and Necrosis Assay are presented as luminescence values relative to the values of DMSO-treated cells at 2 h and plotted as the mean ± s.e.m. (n = 3). *, P < 0.05; **, P < 0.01; ***, P < 0.001, determined by Student’s *t* test using YKL-5-124 and YKL-5-124 + 3-MB-PP1 (F), and YKL-5-124 and YKL-5-124 + iCDK9 (F) data sets.

**Supplementary Table S1A.** Antibodies used in the study.

| Antibody | Source | Identifier |
| --- | --- | --- |
| Mouse monoclonal anti-CDK9 | Santa Cruz Biotechnology | Cat#sc-13130; RRID: AB_627245 |
| Mouse monoclonal anti-p53 | Santa Cruz Biotechnology | Cat#sc-126; RRID: AB_628082 |
| Mouse monoclonal anti-p21 | Santa Cruz Biotechnology | Cat#sc-53870; RRID: AB_785026 |
| Mouse monoclonal anti-GAPDH | Santa Cruz Biotechnology | Cat#sc-32233; RRID: AB_627679 |
| Rabbit polyclonal anti-CDK12 | Cell Signaling Technology | Cat#11973S; RRID: AB_2715688 |
| Goat polyclonal anti-HEXIM1 | Everest Biotech | Cat#EB06964; RRID: AB_2118260 |
| Mouse monoclonal anti-CDK7 | Cell Signaling Technology | Cat#2916; RRID: AB_2077142 |

**Supplementary Table S1B.**
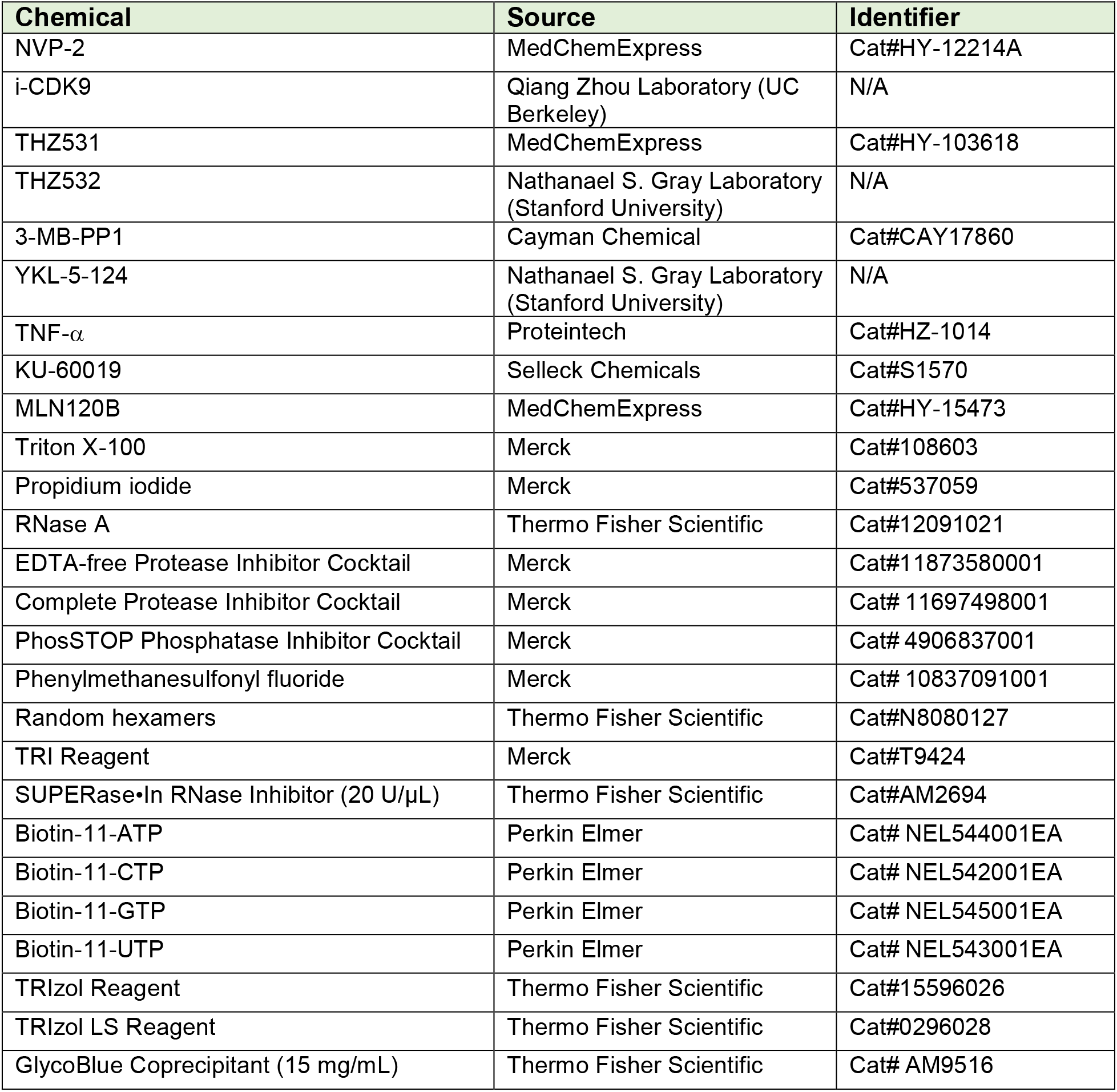
Chemicals used in the study.

| Chemical | Source | Identifier |
| --- | --- | --- |
| NVP-2 | MedChemExpress | Cat#HY-12214A |
| i-CDK9 | Qiang Zhou Laboratory (UC Berkeley) | N/A |
| THZ531 | MedChemExpress | Cat#HY-103618 |
| THZ532 | Nathanael S. Gray Laboratory (Stanford University) | N/A |
| 3-MB-PP1 | Cayman Chemical | Cat#CAY17860 |
| YKL-5-124 | Nathanael S. Gray Laboratory (Stanford University) | N/A |
| TNF-α | Proteintech | Cat#HZ-1014 |
| KU-60019 | Selleck Chemicals | Cat#S1570 |
| MLN120B | MedChemExpress | Cat#HY-15473 |
| Triton X-100 | Merck | Cat#108603 |
| Propidium iodide | Merck | Cat#537059 |
| RNase A | Thermo Fisher Scientific | Cat#12091021 |
| EDTA-free Protease Inhibitor Cocktail | Merck | Cat#11873580001 |
| Complete Protease Inhibitor Cocktail | Merck | Cat# 11697498001 |
| PhosSTOP Phosphatase Inhibitor Cocktail | Merck | Cat# 4906837001 |
| Phenylmethanesulfonyl fluoride | Merck | Cat# 10837091001 |
| Random hexamers | Thermo Fisher Scientific | Cat#N8080127 |
| TRI Reagent | Merck | Cat#T9424 |
| SUPERase•In RNase Inhibitor (20 U/μL) | Thermo Fisher Scientific | Cat#AM2694 |
| Biotin-11-ATP | Perkin Elmer | Cat# NEL544001EA |
| Biotin-11-CTP | Perkin Elmer | Cat# NEL542001EA |
| Biotin-11-GTP | Perkin Elmer | Cat# NEL545001EA |
| Biotin-11-UTP | Perkin Elmer | Cat# NEL543001EA |
| TRIzol Reagent | Thermo Fisher Scientific | Cat#15596026 |
| TRIzol LS Reagent | Thermo Fisher Scientific | Cat#0296028 |
| GlycoBlue Coprecipitant (15 mg/mL) | Thermo Fisher Scientific | Cat# AM9516 |

**Supplementary Table S1C.** Commercial assays used in the study.

| Assay | Source | Identifier |
| --- | --- | --- |
| CellTox™ Green Cytotoxicity Assay | Promega | Cat#G8731 |
| CellTiter-Glo® 2.0 Cell Viability Assay | Promega | Cat#G242 |
| RealTime-Glo Annexin V Apoptosis and Necrosis Assay | Promega | Cat#JA1011 |
| NP-40 lysis buffer | Thermo Fisher Scientific | Cat#J60766-AP |
| M-MLV reverse transcriptase | Thermo Fisher Scientific | Cat#28025-013 |
| Turbo DNA-free™ kit | Thermo Fisher Scientific | Cat#AM1907 |
| Dynabeads Protein G | Thermo Fisher Scientific | Cat#10004D |
| FastStart Universal SYBR Green QPCR Master (Rox) | Merck | Cat#4913914001 |
| MycoplasmaCheck detection kit | Eurofins | Cat#50400400 |
| Bio-Spin P-30 Gel Columns, Tris Buffer | Bio-Rad | Cat# 7326231 |
| Dynabeads MyOne Streptavidin C1 | Thermo Fisher Scientific | Cat#65002 |
| T4 RNA Ligase 1 (ssRNA Ligase) | New England Biolabs | Cat#M0204S |
| RNA 5' Pyrophosphohydrolase (RppH) | New England Biolabs | Cat#M0356S |
| T4 Polynucleotide Kinase | New England Biolabs | Cat# M0201S |
| SuperScript III Reverse Transcriptase | Thermo Fisher Scientific | Cat# 18080044 |
| Phusion Polymerase | In-house | N/A |
| Mag-Bind TotalPure NGS | Omega Bio-tek | Cat# M1378-00 |

**Supplementary Table S1D.**
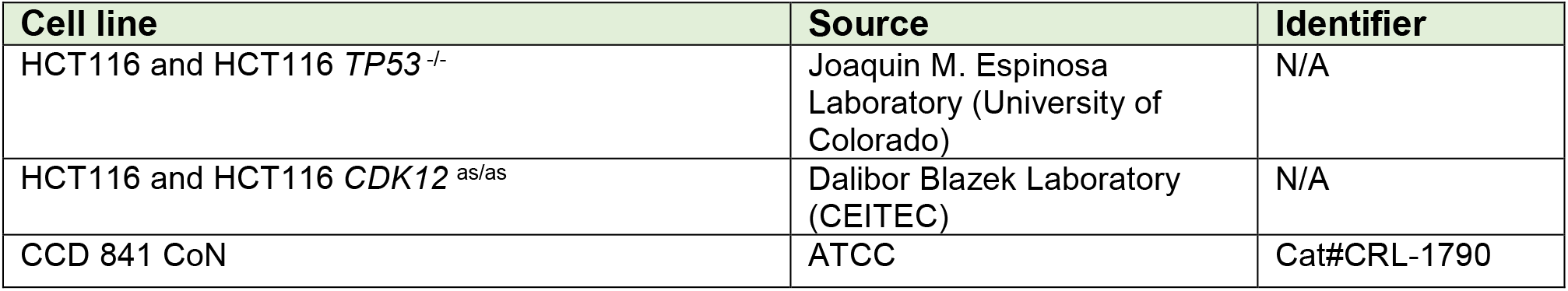
Cell lines used in the study.

**Supplementary Table S1E.** Software and algorithms used in the study.

| Software | Source | Identifier |
| --- | --- | --- |
| R | N/A | https://www.R-project.org/ |
| Molecular Signatures Database v6.0 | GSEA - Broad Institute (Subramanian et al., 2005) | http://software.broadinstitute.org/gsea/msigdb/index.jsp |
| dREG | Wang et al., 2019 | https://dreg.js2.scigap.org |
| Bowtie2 | Langmead & Salzberg, 2012 | https://bowtie-bio.sourceforge.net/bowtie2/index.shtml |
| Samtools | Li et al., 2009 | http://www.htslib.org |
| Fastp | Chen et al., 2018 | https://github.com/OpenGene/fastp |
| Bedtools | Quinlan Laboratory, University of Utah (Quinlan & Hall, 2010) | https://github.com/arq5x/bedtools2 |

**Supplementary Table S1F.**
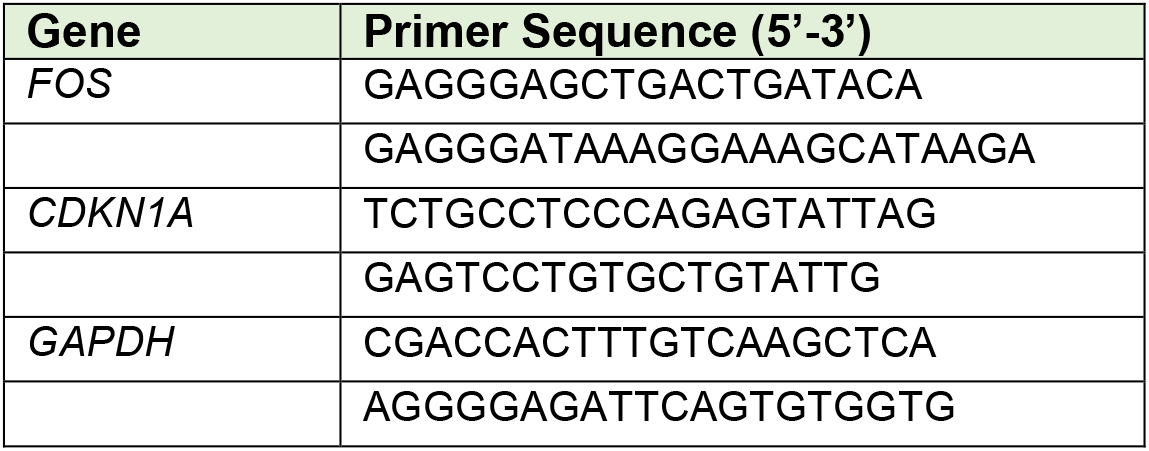
DNA oligonucleotides used in RT-qPCR assay.

**Supplementary Table S1G.**
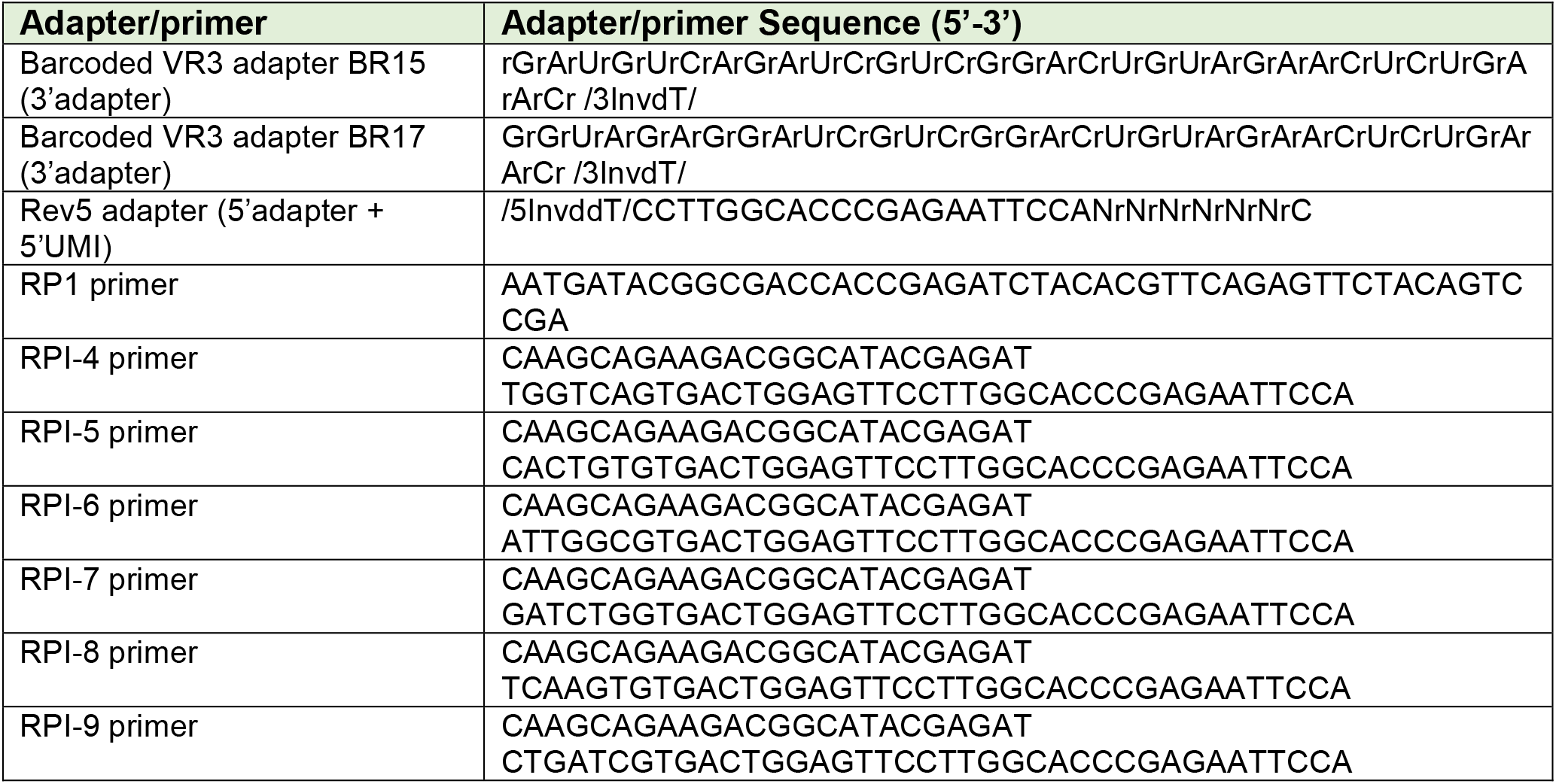
Adapter and primer sequences used in PRO-seq.

**Supplementary Table S2A.**
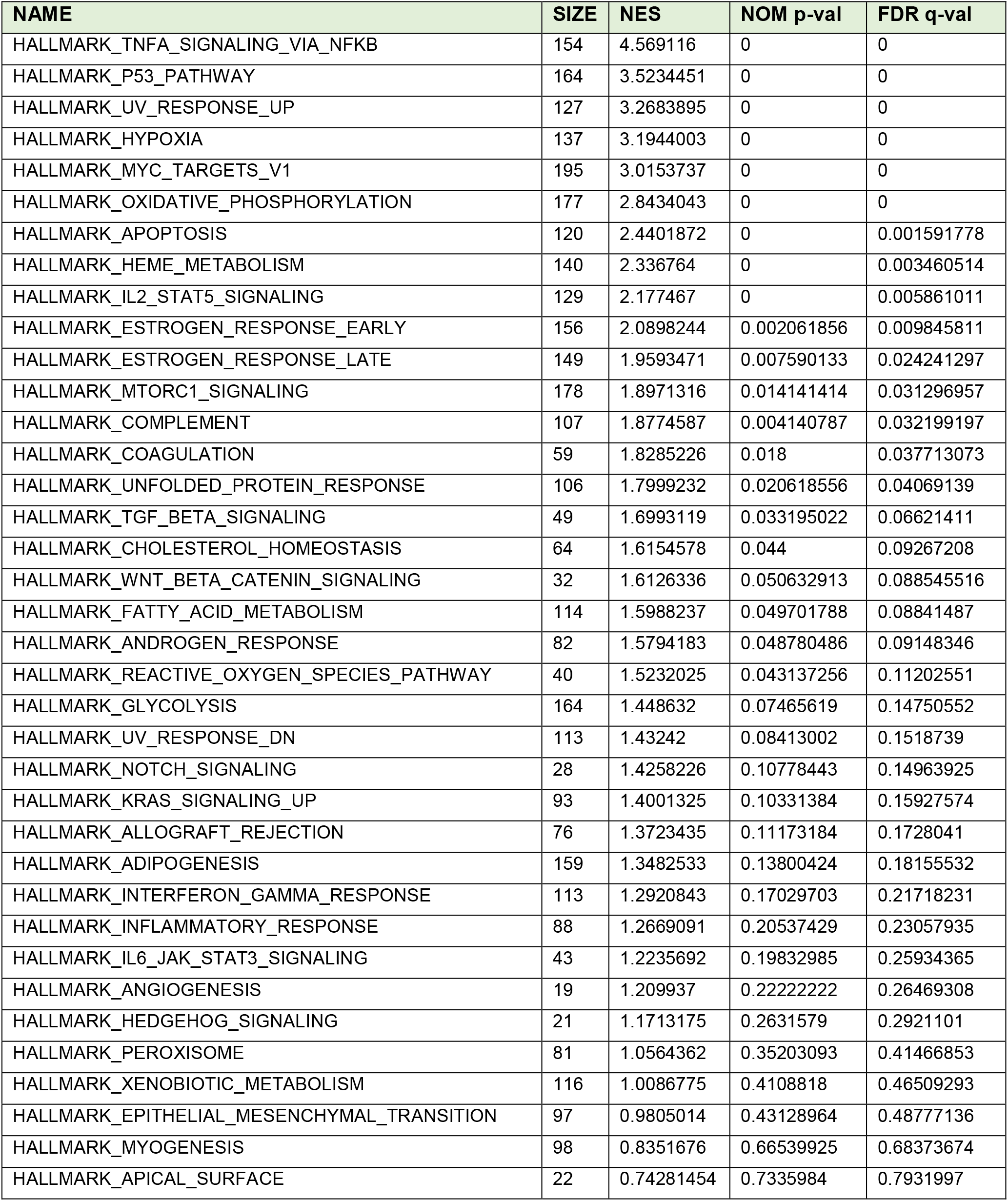
GSEA results of nuclear RNA-seq data from HCT116 CDK12 as/as cell line treated with DMSO or 3-MB-PP1 (5 mM) for 4.5 h. Gene sets enriched among induced genes (NES >0) are shown.

**Supplementary Table S2B.** GSEA results of RNA-seq data from A-375 cell line treated with DMSO or THZ531 (500 nM) for 6 h. Gene sets enriched among induced genes (NES >0) are shown.

| NAME | SIZE | NES | NOM p-val | FDR q-val |
| --- | --- | --- | --- | --- |
| HALLMARK_TNFA_SIGNALING_VIA_NFKB | 169 | 5.9052215 | 0 | 0 |
| HALLMARK_MYC_TARGETS_V1 | 200 | 5.3290915 | 0 | 0 |
| HALLMARK_OXIDATIVE_PHOSPHORYLATION | 199 | 5.269708 | 0 | 0 |
| HALLMARK_MTORC1_SIGNALING | 191 | 3.89095 | 0 | 0 |
| HALLMARK_UV_RESPONSE_UP | 135 | 3.8599348 | 0 | 0 |
| HALLMARK_P53_PATHWAY | 175 | 3.6458375 | 0 | 0 |
| HALLMARK_HYPOXIA | 166 | 3.2016735 | 0 | 0 |
| HALLMARK_ADIPOGENESIS | 181 | 3.0936968 | 0 | 0 |
| HALLMARK_REACTIVE_OXYGEN_SPECIES_PATHWAY | 44 | 2.9286668 | 0 | 0 |
| HALLMARK_FATTY_ACID_METABOLISM | 131 | 2.751583 | 0 | 0 |
| HALLMARK_UNFOLDED_PROTEIN_RESPONSE | 110 | 2.610924 | 0 | 0 |
| HALLMARK_CHOLESTEROL_HOMEOSTASIS | 71 | 2.6005867 | 0 | 0 |
| HALLMARK_XENOBIOTIC_METABOLISM | 135 | 2.5321395 | 0 | 0 |
| HALLMARK_EPITHELIAL_MESENCHYMAL_TRANSITION | 161 | 2.3921928 | 0 | 8.10E-04 |
| HALLMARK_COAGULATION | 81 | 2.3458972 | 0 | 0.00112459 |
| HALLMARK_COMPLEMENT | 134 | 2.3232715 | 0 | 0.001160073 |
| HALLMARK_G2M_CHECKPOINT | 199 | 2.2556317 | 0.003944773 | 0.002461436 |
| HALLMARK_ESTROGEN_RESPONSE_LATE | 146 | 2.2434797 | 0 | 0.002617119 |
| HALLMARK_INFLAMMATORY_RESPONSE | 108 | 2.134734 | 0 | 0.005821647 |
| HALLMARK_APOPTOSIS | 136 | 2.0924578 | 0.00203666 | 0.007158149 |
| HALLMARK_GLYCOLYSIS | 174 | 2.0190966 | 0.002155172 | 0.009531456 |
| HALLMARK_E2F_TARGETS | 200 | 2.0134418 | 0.001964637 | 0.009414393 |
| HALLMARK_HEME_METABOLISM | 161 | 1.931325 | 0.012195122 | 0.01379901 |
| HALLMARK_ANDROGEN_RESPONSE | 89 | 1.8278781 | 0.003976143 | 0.025065133 |
| HALLMARK_ALLOGRAFT_REJECTION | 103 | 1.814645 | 0.014112903 | 0.025645318 |
| HALLMARK_DNA_REPAIR | 146 | 1.6505222 | 0.029880479 | 0.058037277 |
| HALLMARK_ANGIOGENESIS | 22 | 1.5723708 | 0.058479533 | 0.08264521 |
| HALLMARK_IL2_STAT5_SIGNALING | 147 | 1.5504019 | 0.049701788 | 0.088566504 |
| HALLMARK_PANCREAS_BETA_CELLS | 17 | 1.5439894 | 0.04897959 | 0.08847765 |
| HALLMARK_APICAL_JUNCTION | 141 | 1.5326812 | 0.06313646 | 0.08998745 |
| HALLMARK_PROTEIN_SECRETION | 92 | 1.5123383 | 0.07602339 | 0.095925584 |
| HALLMARK_MYC_TARGETS_V2 | 58 | 1.4882184 | 0.0804829 | 0.1018874 |
| HALLMARK_MYOGENESIS | 128 | 1.4422815 | 0.09594883 | 0.120485745 |
| HALLMARK_TGF_BETA_SIGNALING | 50 | 1.41272 | 0.106299214 | 0.13244651 |
| HALLMARK_PEROXISOME | 83 | 1.3349255 | 0.1364562 | 0.17507945 |
| HALLMARK_WNT_BETA_CATENIN_SIGNALING | 31 | 1.2701924 | 0.17226891 | 0.21925417 |
| HALLMARK_PI3K_AKT_MTOR_SIGNALING | 89 | 1.2146652 | 0.21010101 | 0.26125103 |
| HALLMARK_INTERFERON_GAMMA_RESPONSE | 144 | 1.2036312 | 0.21548118 | 0.26381475 |
| HALLMARK_SPERMATOGENESIS | 74 | 1.1934415 | 0.25851703 | 0.26638502 |
| HALLMARK_BILE_ACID_METABOLISM | 65 | 1.0461472 | 0.38011697 | 0.41898945 |
| HALLMARK_UV_RESPONSE_DN | 132 | 1.0401976 | 0.398 | 0.4159044 |
| HALLMARK_INTERFERON_ALPHA_RESPONSE | 79 | 0.75348884 | 0.766 | 0.81790626 |
| HALLMARK_NOTCH_SIGNALING | 29 | 0.7158327 | 0.8426295 | 0.8480125 |
| HALLMARK_ESTROGEN_RESPONSE_EARLY | 160 | 0.7078182 | 0.8359375 | 0.8377617 |

**Supplementary Table S2C.** GSEA results of RNA-seq data from Colo829 cell line treated with DMSO or THZ531 (500 nM) for 6 h. Gene sets enriched among induced genes (NES >0) are shown.

| NAME | SIZE | NES | NOM p-val | FDR q-val |
| --- | --- | --- | --- | --- |
| HALLMARK_TNFA_SIGNALING_VIA_NFKB | 170 | 5.993585 | 0 | 0 |
| HALLMARK_MYC_TARGETS_V1 | 200 | 4.8723793 | 0 | 0 |
| HALLMARK_EPITHELIAL_MESENCHYMAL_TRANSITION | 159 | 4.440328 | 0 | 0 |
| HALLMARK_OXIDATIVE_PHOSPHORYLATION | 200 | 4.3971496 | 0 | 0 |
| HALLMARK_MTORC1_SIGNALING | 191 | 3.7962265 | 0 | 0 |
| HALLMARK_HYPOXIA | 170 | 3.765153 | 0 | 0 |
| HALLMARK_P53_PATHWAY | 174 | 3.6782324 | 0 | 0 |
| HALLMARK_UV_RESPONSE_UP | 135 | 3.2540934 | 0 | 0 |
| HALLMARK_APOPTOSIS | 133 | 2.7054396 | 0 | 3.37E-04 |
| HALLMARK_REACTIVE_OXYGEN_SPECIES_PATHWAY | 43 | 2.683158 | 0 | 3.03E-04 |
| HALLMARK_COAGULATION | 87 | 2.4981923 | 0 | 4.67E-04 |
| HALLMARK_COMPLEMENT | 144 | 2.446013 | 0 | 8.10E-04 |
| HALLMARK_IL2_STAT5_SIGNALING | 151 | 2.264054 | 0 | 0.003182184 |
| HALLMARK_GLYCOLYSIS | 178 | 2.1689503 | 0 | 0.005267459 |
| HALLMARK_INFLAMMATORY_RESPONSE | 118 | 2.1503437 | 0.002087683 | 0.005576605 |
| HALLMARK_G2M_CHECKPOINT | 199 | 2.114998 | 0 | 0.006373445 |
| HALLMARK_UNFOLDED_PROTEIN_RESPONSE | 110 | 2.0996726 | 0.002066116 | 0.006678859 |
| HALLMARK_ADIPOGENESIS | 182 | 2.0222383 | 0.001908397 | 0.012223416 |
| HALLMARK_CHOLESTEROL_HOMEOSTASIS | 69 | 1.98417 | 0.005964215 | 0.014527635 |
| HALLMARK_XENOBIOTIC_METABOLISM | 130 | 1.8821785 | 0.009363296 | 0.02217932 |
| HALLMARK_TGF_BETA_SIGNALING | 50 | 1.7924047 | 0.01171875 | 0.033797543 |
| HALLMARK_MYC_TARGETS_V2 | 58 | 1.778902 | 0.015936255 | 0.034482583 |
| HALLMARK_HEME_METABOLISM | 164 | 1.772658 | 0.014373717 | 0.034299865 |
| HALLMARK_FATTY_ACID_METABOLISM | 131 | 1.7465398 | 0.034026466 | 0.03743994 |
| HALLMARK_ANGIOGENESIS | 23 | 1.7208741 | 0.03515625 | 0.040528085 |
| HALLMARK_E2F_TARGETS | 200 | 1.6960897 | 0.025896415 | 0.043979794 |
| HALLMARK_ESTROGEN_RESPONSE_LATE | 149 | 1.6078511 | 0.029469548 | 0.0633915 |
| HALLMARK_WNT_BETA_CATENIN_SIGNALING | 35 | 1.5616962 | 0.049523808 | 0.07616374 |
| HALLMARK_DNA_REPAIR | 146 | 1.5555179 | 0.061420344 | 0.07576268 |
| HALLMARK_ANDROGEN_RESPONSE | 92 | 1.5367256 | 0.049079753 | 0.08001442 |
| HALLMARK_SPERMATOGENESIS | 75 | 1.4359711 | 0.09356725 | 0.12506118 |
| HALLMARK_APICAL_JUNCTION | 141 | 1.3053916 | 0.16796875 | 0.2082934 |
| HALLMARK_ALLOGRAFT_REJECTION | 106 | 1.2827123 | 0.18476191 | 0.21930678 |
| HALLMARK_MYOGENESIS | 138 | 1.261024 | 0.18571429 | 0.22950968 |
| HALLMARK_PI3K_AKT_MTOR_SIGNALING | 87 | 1.2046958 | 0.21643287 | 0.27308276 |
| HALLMARK_UV_RESPONSE_DN | 138 | 1.2012941 | 0.22954091 | 0.26856044 |
| HALLMARK_MITOTIC_SPINDLE | 198 | 1.120099 | 0.2835821 | 0.34163022 |
| HALLMARK_PEROXISOME | 84 | 1.1015757 | 0.3138833 | 0.3524855 |
| HALLMARK_KRAS_SIGNALING_UP | 121 | 1.0728679 | 0.34738955 | 0.37429097 |
| HALLMARK_HEDGEHOG_SIGNALING | 28 | 0.9446006 | 0.4990403 | 0.5341227 |
| HALLMARK_NOTCH_SIGNALING | 28 | 0.7393361 | 0.8041667 | 0.8214937 |
| HALLMARK_IL6_JAK_STAT3_SIGNALING | 55 | 0.7217652 | 0.8040816 | 0.824549 |

**Supplementary Table S2D.** GSEA results of RNA-seq data from MDA-MB-231 cell line treated with DMSO or SR-4835 (90 nM) for 6 h. Gene sets enriched among induced genes (NES >0) are shown.

| NAME | SIZE | NES | NOM p-val | FDR q-val |
| --- | --- | --- | --- | --- |
| HALLMARK_MYC_TARGETS_V1 | 195 | 4.2677917 | 0 | 0 |
| HALLMARK_TNFA_SIGNALING_VIA_NFKB | 195 | 3.4495616 | 0 | 0 |
| HALLMARK_OXIDATIVE_PHOSPHORYLATION | 185 | 3.2095857 | 0 | 0 |
| HALLMARK_MTORC1_SIGNALING | 196 | 2.8713398 | 0 | 0 |
| HALLMARK_UV_RESPONSE_UP | 149 | 2.6897907 | 0 | 3.10E-04 |
| HALLMARK_HYPOXIA | 191 | 2.343713 | 0 | 0.001378287 |
| HALLMARK_REACTIVE_OXYGEN_SPECIES_PATHWAY | 48 | 2.0940063 | 0.001996008 | 0.00812981 |
| HALLMARK_UNFOLDED_PROTEIN_RESPONSE | 109 | 2.085606 | 0.001976285 | 0.007288878 |
| HALLMARK_ESTROGEN_RESPONSE_LATE | 187 | 1.706714 | 0.011976048 | 0.04967046 |
| HALLMARK_P53_PATHWAY | 190 | 1.6474636 | 0.035856575 | 0.059560914 |
| HALLMARK_CHOLESTEROL_HOMEOSTASIS | 71 | 1.4680228 | 0.11 | 0.12338756 |
| HALLMARK_PANCREAS_BETA_CELLS | 27 | 1.4061323 | 0.10748561 | 0.14729646 |
| HALLMARK_ANDROGEN_RESPONSE | 96 | 1.3552537 | 0.10891089 | 0.16598897 |
| HALLMARK_PI3K_AKT_MTOR_SIGNALING | 104 | 1.351604 | 0.1211499 | 0.15605791 |
| HALLMARK_ALLOGRAFT_REJECTION | 169 | 1.344997 | 0.120162934 | 0.15006815 |
| HALLMARK_HEDGEHOG_SIGNALING | 35 | 0.7825379 | 0.73333335 | 0.78388757 |
| HALLMARK_ANGIOGENESIS | 35 | 0.65965456 | 0.8816568 | 0.8898483 |

**Supplementary Table S2E.** GSEA results of TT-seq data from IMR-32 cell line treated with DMSO or THZ531 (400 nM) for 2 h. Gene sets enriched among induced genes (NES >0) are shown.

| NAME | SIZE | NES | NOM p-val | FDR q-val |
| --- | --- | --- | --- | --- |
| HALLMARK_MYC_TARGETS_V1 | 193 | 5.563506 | 0 | 0 |
| HALLMARK_OXIDATIVE_PHOSPHORYLATION | 177 | 5.0663924 | 0 | 0 |
| HALLMARK_TNFA_SIGNALING_VIA_NFKB | 99 | 4.719935 | 0 | 0 |
| HALLMARK_UV_RESPONSE_UP | 112 | 4.197955 | 0 | 0 |
| HALLMARK_P53_PATHWAY | 125 | 3.3015583 | 0 | 0 |
| HALLMARK_MTORC1_SIGNALING | 183 | 3.092014 | 0 | 0 |
| HALLMARK_DNA_REPAIR | 135 | 3.0866828 | 0 | 0 |
| HALLMARK_HYPOXIA | 127 | 2.9120805 | 0 | 0 |
| HALLMARK_XENOBIOTIC_METABOLISM | 95 | 2.8804173 | 0 | 0 |
| HALLMARK_APOPTOSIS | 98 | 2.8326726 | 0 | 0 |
| HALLMARK_UNFOLDED_PROTEIN_RESPONSE | 102 | 2.5896707 | 0 | 2.99E-04 |
| HALLMARK_REACTIVE_OXYGEN_SPECIES_PATHWAY | 40 | 2.549103 | 0 | 4.76E-04 |
| HALLMARK_FATTY_ACID_METABOLISM | 108 | 2.5357153 | 0.002024292 | 4.39E-04 |
| HALLMARK_ADIPOGENESIS | 147 | 2.5286474 | 0 | 4.08E-04 |
| HALLMARK_MYC_TARGETS_V2 | 54 | 2.3626895 | 0 | 0.00146277 |
| HALLMARK_PANCREAS_BETA_CELLS | 15 | 2.232777 | 0.001923077 | 0.003968566 |
| HALLMARK_PEROXISOME | 74 | 2.171232 | 0.001996008 | 0.005409245 |
| HALLMARK_E2F_TARGETS | 195 | 2.1331124 | 0.001960784 | 0.00593865 |
| HALLMARK_MYOGENESIS | 79 | 2.0512328 | 0.007874016 | 0.008701 |
| HALLMARK_GLYCOLYSIS | 140 | 1.9735106 | 0.004 | 0.013776437 |
| HALLMARK_INTERFERON_GAMMA_RESPONSE | 79 | 1.9194916 | 0.015209125 | 0.017592223 |
| HALLMARK_EPITHELIAL_MESENCHYMAL_TRANSITION | 92 | 1.8656666 | 0.011976048 | 0.023216793 |
| HALLMARK_G2M_CHECKPOINT | 185 | 1.855756 | 0.00952381 | 0.023416903 |
| HALLMARK_ESTROGEN_RESPONSE_LATE | 109 | 1.8553507 | 0.015122874 | 0.0226012 |
| HALLMARK_COAGULATION | 39 | 1.844479 | 0.015686275 | 0.023018748 |
| HALLMARK_ALLOGRAFT_REJECTION | 59 | 1.76368 | 0.019417476 | 0.03456152 |
| HALLMARK_APICAL_JUNCTION | 94 | 1.7442787 | 0.02 | 0.036381908 |
| HALLMARK_COMPLEMENT | 92 | 1.7105255 | 0.025917927 | 0.041899767 |
| HALLMARK_KRAS_SIGNALING_UP | 72 | 1.6993793 | 0.03168317 | 0.04257905 |
| HALLMARK_CHOLESTEROL_HOMEOSTASIS | 57 | 1.6798801 | 0.04474708 | 0.04570205 |
| HALLMARK_IL6_JAK_STAT3_SIGNALING | 30 | 1.5345051 | 0.068359375 | 0.08708852 |
| HALLMARK_NOTCH_SIGNALING | 20 | 1.5226976 | 0.052941177 | 0.08898578 |
| HALLMARK_ESTROGEN_RESPONSE_EARLY | 109 | 1.4888661 | 0.09259259 | 0.10064545 |
| HALLMARK_PI3K_AKT_MTOR_SIGNALING | 80 | 1.3742985 | 0.1254902 | 0.16319123 |
| HALLMARK_BILE_ACID_METABOLISM | 52 | 1.3338807 | 0.13229571 | 0.1839347 |
| HALLMARK_WNT_BETA_CATENIN_SIGNALING | 27 | 1.2687906 | 0.1743487 | 0.23063491 |
| HALLMARK_ANDROGEN_RESPONSE | 76 | 1.2359577 | 0.21869159 | 0.2526209 |
| HALLMARK_HEME_METABOLISM | 125 | 1.2334836 | 0.1987705 | 0.2480089 |
| HALLMARK_IL2_STAT5_SIGNALING | 103 | 1.2308427 | 0.21399178 | 0.24436803 |
| HALLMARK_INTERFERON_ALPHA_RESPONSE | 35 | 1.1718471 | 0.2637131 | 0.2934957 |
| HALLMARK_TGF_BETA_SIGNALING | 43 | 1.169077 | 0.24950495 | 0.2889714 |
| HALLMARK_KRAS_SIGNALING_DN | 45 | 1.1259705 | 0.30737704 | 0.3246291 |
| HALLMARK_SPERMATOGENESIS | 66 | 1.1041087 | 0.32629558 | 0.341718 |
| HALLMARK_INFLAMMATORY_RESPONSE | 50 | 1.0801312 | 0.35546038 | 0.35893694 |
| HALLMARK_ANGIOGENESIS | 15 | 0.8187178 | 0.6785714 | 0.70232785 |
| HALLMARK_HEDGEHOG_SIGNALING | 26 | 0.68658835 | 0.87090164 | 0.8625199 |

